# Foreign body reaction is triggered *in vivo* by cellular mechanosensing of implants stiffer than host tissue

**DOI:** 10.1101/829648

**Authors:** Alejandro Carnicer-Lombarte, Damiano G. Barone, Ivan B. Dimov, Russell S. Hamilton, Malwina Prater, Xiaohui Zhao, Alexandra L. Rutz, George G. Malliaras, Stephanie P. Lacour, Clare E. Bryant, James W. Fawcett, Kristian Franze

**Affiliations:** John van Geest Centre for Brain Repair, Department of Clinical Neurosciences, University of Cambridge, Cambridge CB2 0PY, UK; Department of Physiology Development and Neuroscience, University of Cambridge, Cambridge CB2 3DY, UK; Department of Veterinary Medicine, University of Cambridge, Madingley Road, CambridgèCB3 0ES, UK; Centre for Trophoblast Research, University of Cambridge, Cambridge, CB2 3EG, UK; Electrical Engineering Division, Department of Engineering, University of Cambridge, Cambridge CB3 0FA, UK; Bertarelli Foundation Chair in Neuroprosthetic Technology, Laboratory for Soft Bioelectronics Interface, Institute of Microengineering, Institute of Bioengineering, Centre for Neuroprosthetics, Ecole Polytechnique Fédérale de Lausanne (EPFL), 1202 Geneva, Switzerland; Centre for Reconstructive Neuroscience, Institute for Experimental Medicine CAS, Prague, Czech Republic; Institute of Medical Physics, Friedrich-Alexander University Erlangen-Nuremberg, 91052 Erlangen, Germany; Max-Planck-Zentrum für Physik und Medizin, 91054 Erlangen, Germany

## Abstract

Medical implants offer a unique and powerful therapeutic approach in many areas of medicine. However, their lifetime is often limited as they may cause a foreign body reaction (FBR) leading to their encapsulation by scar tissue^1–4^. Despite the importance of this process, how cells recognise implanted materials is still poorly understood^5, 6^.

Here, we show how the mechanical mismatch between implants and host tissue leads to FBR. Fibroblasts and macrophages, which are both crucially involved in mediating FBR, became activated when cultured on materials just above the stiffness of healthy tissue. Coating stiff implants with a thin layer of hydrogel or silicone with a tissue-like elastic modulus (∼20 kPa in subcutaneous and ∼2 kPa in peripheral nerve implants) or softer significantly reduced inflammation and fibrosis three months after implantation. Materials stiffer than the host tissue led to nuclear localisation of the mechanosensitive transcriptional regulator YAP in neighbouring cells *in vivo*, confirming mechanotransduction. The alleviation of FBR by soft coatings not exceeding the stiffness of the host tissue provides a strategy to achieve long-term implant stability without extensive modification of current implant manufacturing techniques, facilitating clinical translation.

## Introduction

Medical implants have become an indispensable tool for a wide range of applications, including bladder control^7^, treatment of neurological disorders^8^, drug delivery^9^, and tissue repair^10^. Recent advances in fabrication techniques have allowed for the development of increasingly complex implants, capable of better integrating into the host tissue and much improved functionality. This is particularly visible in the field of neural interfaces, where implants are capable of establishing electrical connections with individual axons and neurons without disrupting their connectivity^11, 12^. Functionality of such implants actively interacting with their environment requires the establishment and maintenance of intimate interfaces between implant and tissue.

However, this interface and thus long-term functionality of medical implants is often limited by foreign body reaction (FBR) – a process by which the body recognises implanted materials as foreign and attempts to degrade them. FBR is characterised by chronic inflammation and fibrosis, and with time results in the formation of a scar – a fibrotic capsule – separating the implant from the host tissue^5^. How the body recognises implanted materials as foreign and what triggers the associated inflammatory response is still not fully understood^5, 6^.

The breakdown of the tissue-implant interface is one of the leading causes of implant failure^1–3^, which has led to large efforts to develop a treatment to prevent and manage FBR to implanted materials^13, 14^. Impregnation of implants with anti-inflammatory drugs such as dexamethasone is currently widely used in clinical practice to alleviate inflammatory reactions and suppress FBR in nerve interfaces^15^. However, such chemical treatments can have significant side effects^15^.

Recent studies indicated that softer materials may also alleviate FBR^16–19^. However, to date we do not know how soft materials need to be in order to avoid foreign body reactions. In fact, elastic moduli (a measure of stiffness) of implant materials classified as ‘soft’ in literature range from kPa to GPa – spanning 6 orders of magnitude^20^. Here we show that implant materials need to be at least as soft as the host tissue to suppress FBR. We show that FBR is triggered by cellular mechanotransduction at the surface of stiff implants *in vivo* rather than by tissue damage due to rigid implants as often thought^20–22^. Furthermore, we developed a scalable technique to manufacture implants with a thin layer of a very soft material – down to the softness of the host tissue itself – which accommodates cellular mechanosensitivity and greatly reduces FBR.

## Results

We first investigated how primary macrophages and fibroblasts, cell types crucial for FBR in many tissue types, respond to substrates with different mechanical properties, covering a physiologically relevant stiffness range. Macrophages are a central component of innate immunity and are responsible for driving the inflammatory response to implanted materials^5, 6^. Fibroblasts, on the other hand, are the primary mediators of the fibrotic response, responsible for the formation of the fibrotic capsule around the implant.

Most macrophages involved in FBR are derived from blood-circulating monocytes, while fibroblasts proliferate from tissue-resident populations^5, 6^. We cultured primary bone marrow- derived macrophages and fibroblast populations derived from peripheral nerve tissue on polyacrylamide substrates of a range of elastic (shear) moduli. We chose nerve fibroblasts because of the high impact of FBR on peripheral nerve interfaces. While implants with shear moduli of few hundreds of kPa are commonly considered ‘soft’^21–23^, shear moduli of our substrates ranged from 0.1 kPa, mechanically resembling the softest tissues in the body including peripheral nerve tissue^24^, to 50 kPa, which is already stiffer than most soft tissues and organs^25^ (substrates used: 0.1, 1, 10, 50 kPa shear modulus; corresponding to ∼0.3, 3, 30, 150 kPa Young’s modulus, Supplementary Fig. 1).

FBR-induced scarring is particularly detrimental to electrical neural interfaces, such as those implanted in nerves. We therefore initially focused on this tissue. Nerve fibroblasts cultured on substrates of varying stiffness for 6 days retained a spherical morphology on soft gels with shear moduli of ∼0.1-1 kPa, while cell spreading significantly increased on stiffer substrates with shear moduli of 10-50 kPa (Fig. 1b) (*p* = 1.2e-7, one-way ANOVA). Adhesion to the substrate via focal adhesions (Fig. 1c) and proliferation (Supplementary Fig. 2) showed similar significant increases with substrate stiffness (adhesion *p* = 5.2e-9, proliferation *p* = 0.0023). These observed increases in cell spreading, adhesion, and proliferation are consistent with a fibroblast FBR-like phenotype^26^.

**Fig 1.**
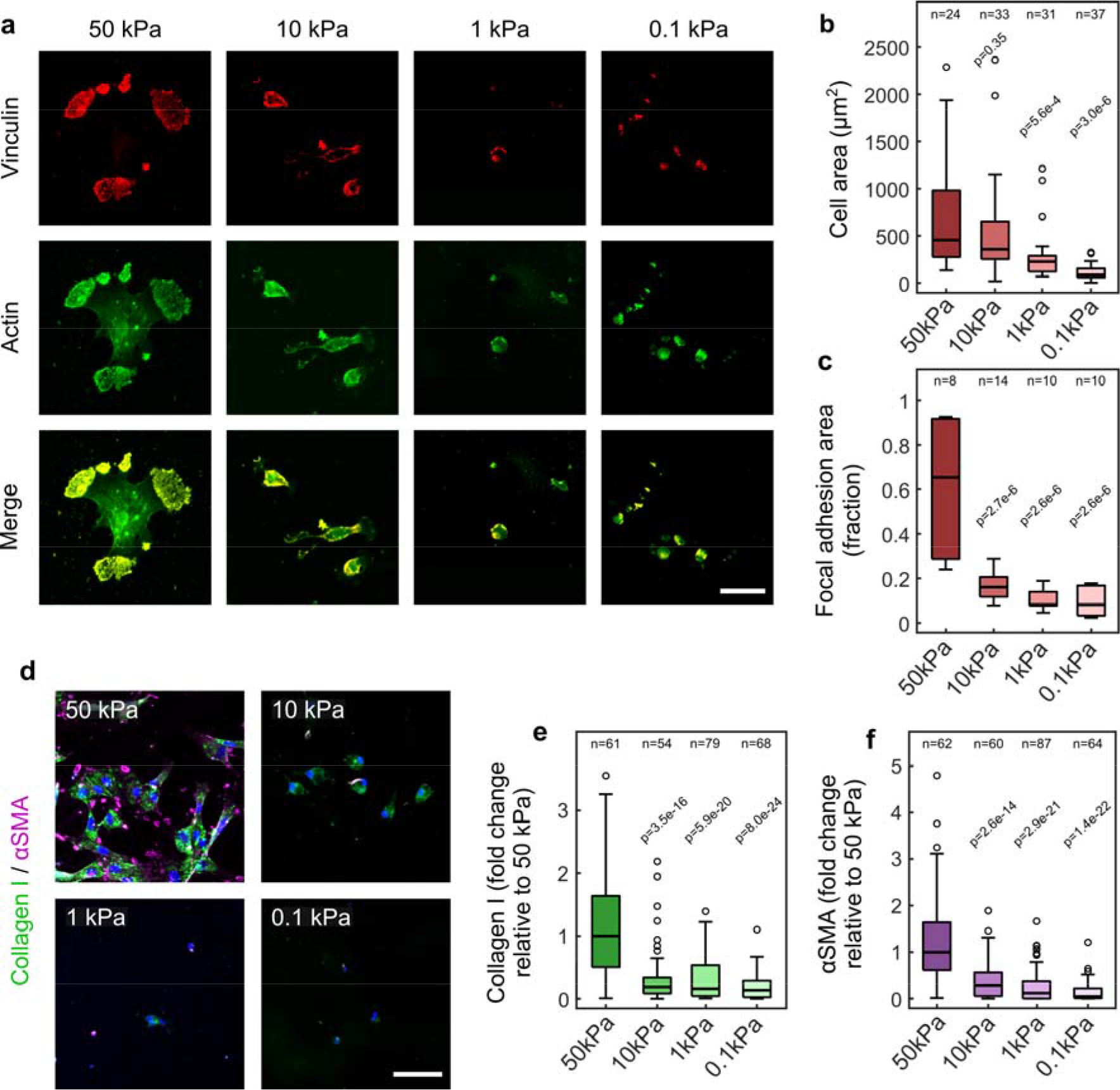
Substrates with a stiffness above that of tissue trigger fibrosis *in vitro*. **a**, Maximum intensity projections of *z*-stack confocal images of nerve fibroblasts at 6 DIV (days *in vitro*) cultured on polyacrylamide substrates of various stiffness (50, 10, 1 and 0.1 kPa shear modulus), stained for cytoskeleton (actin) and focal adhesion (vinculin) markers. Cell morphologies significantly changed on non-physiologically stiff (shear modulus of 50 kPa) substrates. Scale bar: 30 μm. **b**,**c**, Box plots of fibroblast cell area (**b**) and focal adhesion area (**c**). *n* = number of cells. **d**, Images of fibroblasts stained for myofibroblast markers αSMA (magenta) and collagen I (green) show an increase in fibrotic phenotype on 50 kPa gels. Cell nuclei stained with DAPI (blue) Scale bar: 60 μm. **e**,**f**, Box plots of relative stain intensities for collagen I (**e**) and αSMA (**f**). *n* = number of cells. Box plot characteristics described in Methods. All statistical comparisons done via one-way ANOVA followed by Dunnett’s multiple comparisons test comparing to the 50 kPa condition. All experiments performed 3 - 4 times.

Fibroblasts in FBR and other fibrotic processes typically differentiate into myofibroblasts – which are characterised by the expression of alpha smooth muscle actin (αSMA) and high extracellular matrix production^27^. Exposure to the non-physiological high stiffness of 50 kPa significantly increased the synthesis of both αSMA and the extracellular matrix protein collagen I – a primary component of the FBR capsule (Fig. 1d-f) (collagen *p* = 1.4e-26, αSMA *p* = 5.3e-26, one-way ANOVAs). Fibronectin - a different extracellular matrix protein – was present in nerve fibroblast cultures but was largely independent of substrate stiffness (Supplementary Fig. 2). Our results on primary nerve fibroblasts are consistent with previous studies of fibroblast populations from other tissues reporting that fibroblasts transition into a myofibroblast phenotype at high substrate stiffnesses^27, 28^.

Macrophages showed similar functional changes on substrates stiffer than their physiological host tissue. Similar to nerve fibroblasts, cell spreading, adhesion, and proliferation rates were significantly increased on stiffer if compared to softer substrates (Fig 2a-c, Supplementary Fig. 3) (spreading *p* = 9.8e-8, adhesion *p* = 0.009, proliferation *p* = 0.007, one-way ANOVA), indicating macrophage activation.

**Fig 2.**
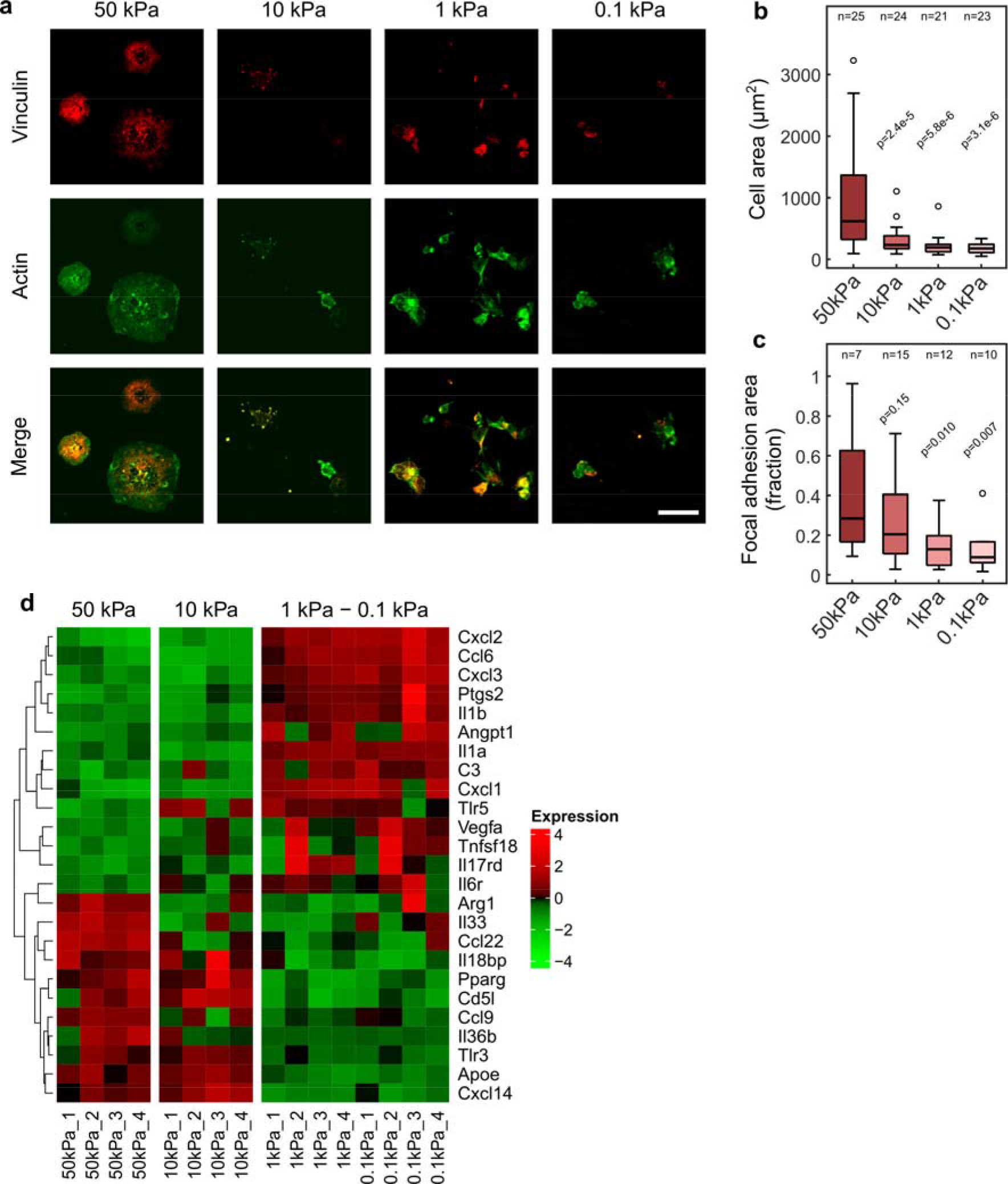
Substrates with a stiffness above that of native tissue trigger macrophage activation and changes in inflammatory profile *in vitro*. a, Maximum intensity projections of *z*-stack confocal images of bone marrow-derived macrophages at 6 DIV (days *in vitro*) cultured on polyacrylamide substrates of various stiffness (50, 10, 1 and 0.1 kPa shear modulus), stained for cytoskeleton (actin) and focal adhesion (vinculin) markers. Cell morphologies significantly changed on non-physiologically stiff (50 kPa) substrates. Scale bar: 30 μm. b,c, Boxplots of macrophage cell area (b) and focal adhesion area (c). *n* = number of cells. All statistical comparisons done via one-way ANOVA followed by Dunnett’s multiple comparisons test comparing to the 50 kPa condition. Box plot characteristics described in Methods. d, Heatmap of changes in inflammatory differentially expressed gene (DEG) expression profile of macrophages at 3 DIV detected in RNAseq (expression values represented as base-2 log fold change of counts, with respect to the average across all groups, as previously used^65^). Changes in markers such as *arg1, pparg, il1b,* and *ptgs2* indicate a switch to an M2-like scar production-driving phenotype of macrophages grown on high stiffness substrates. Four samples analysed for each substrate stiffness condition.

RNA sequencing of macrophages three days post-plating showed a switch in gene expression patterns, particularly in inflammation-related genes, between cells cultured on substrates as soft as or softer than healthy nerve tissue (0.1 – 1 kPa) and cells cultured on substrates stiffer than nerves (10 – 50 kPa) (Fig. 2d). These transcriptional differences were consistent with a switch in phenotype from pro-inflammatory M1 on softer substrates to the scar production-driving M2 phenotype on stiffer substrates. On stiff substrates, we found downregulation of proinflammatory genes such as interleukin 1 beta (*il1b*) (*p* = 1.4e-6, FDR adjusted p-value) and prostaglandin E synthase (*ptgs2*) (*p* = 9.5e-6), while M2-associated genes such as arginase (*arg1*) (*p* = 0.002) and peroxisome proliferator-activated receptor-γ (*ppar*γ) (*p* = 0.007) were upregulated if compared to soft substrates. This switch to a macrophage M2 activation profile is typically associated with tissue regeneration and fibrosis^29, 30^. Together, our *in vitro* experiments indicated that a substrate stiffness above that of the host tissue may lead to fibrosis, a hallmark of FBR.

Having confirmed that both primary macrophages and nerve fibroblasts assume an FBR-like phenotype when exposed to materials that are stiffer than their host tissues, we sought to test if materials mechanically matched to these tissues may alleviate FBR at the cellular level *in vivo* in rat models. While it is often difficult or impractical to fabricate and use implants made entirely from extremely soft materials because of fabrication and handling challenges, stiff materials can be masked from cells underneath a thick enough layer of a softer material^31–33^. Hence, we designed silicone rubber implants with a shear modulus of ∼ 200 kPa, which we coated with a 100 μm thick layer of a soft material (Fig. 3a). The structural mechanical properties of these implants, such as bending stiffness, are dominated by the much thicker ∼200 kPa silicone rubber and are thus largely independent of the coatings. However, at the cellular level, the thin coatings will “stealthen” the underlying stiffer substrate from the adjacent cells (see Methods for details)^31–33^.

**Fig 3.**
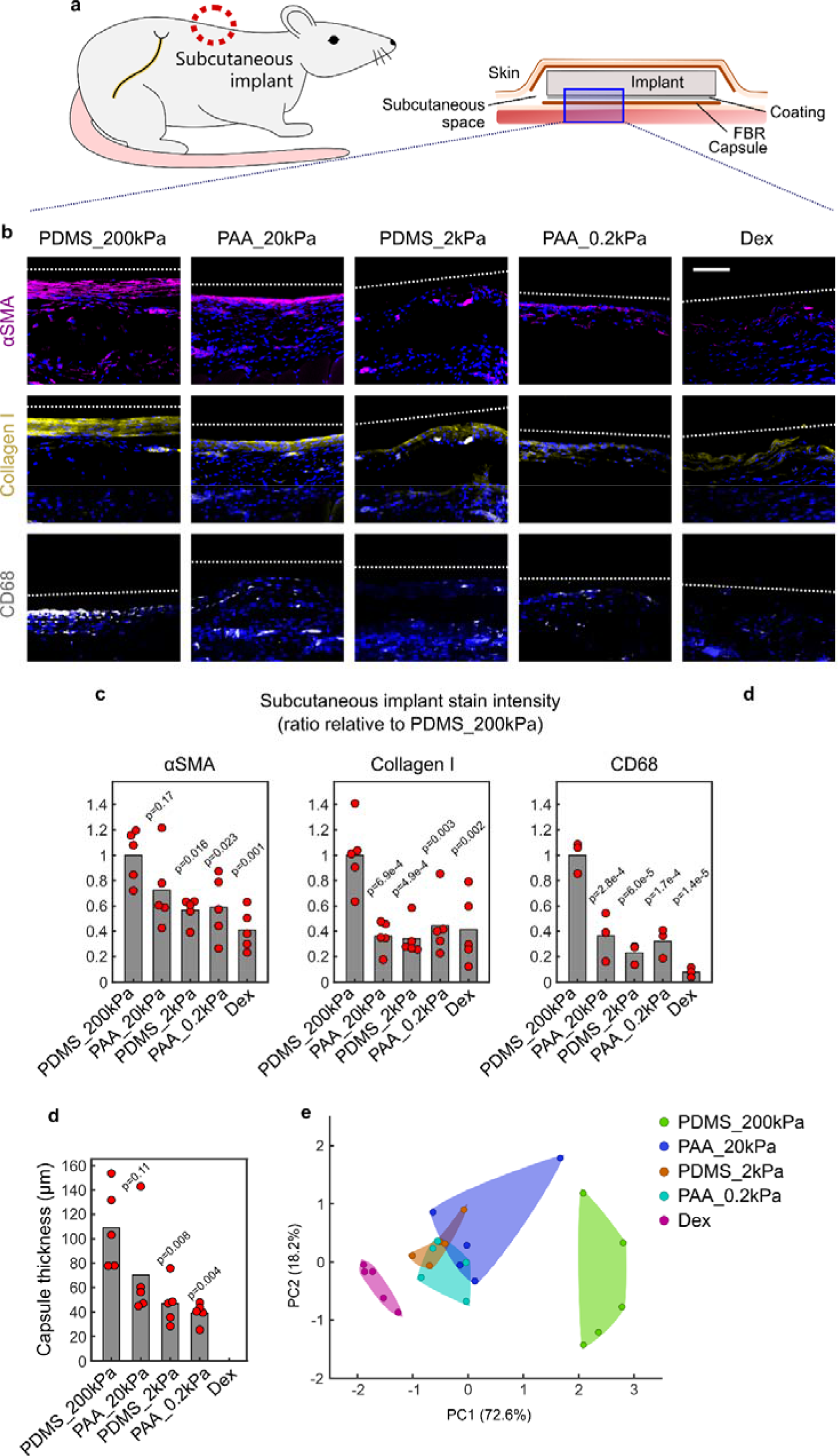
Coatings at least as soft as the host tissue significantly reduce foreign body reaction to subcutaneous implants 3 months post-implantation *in vivo*. a, Diagram of implant location and structure. Blue square represents area shown in images in b. b, *Z*-stack confocal images of subcutaneous tissue in response to implants of 200 kPa shear modulus silicone PDMS (PDMS_200kPa), or to implants coated with 20 kPa polyacrylamide hydrogel (PAA_20kPa), 2 kPa PDMS (PDMS_2kPa), 0.2 kPa polyacrylamide (PAA_0.2kPa), or PDMS impregnated with 10 mg/ml of the anti-inflammatory drug dexamethasone (Dex). Tissue is fluorescently labelled for myofibroblasts (αSMA, magenta), extracellular matrix components (collagen I, yellow), and macrophages (CD68, white). Approximate edges of implant are indicated by dashed white lines. All images show DAPI stains of nuclei (blue). Scale bar: 100 μm. c, Plot of stain intensities. d, Plot of fibrotic capsule thickness. Quantification of thickness in Dex group absent as dexamethasone impeded formation of a structured boundary between tissue and implant. For all plots: bars represent mean, dots represent individual animals. *N* = 5 rats. Statistical comparisons carried out via one-way ANOVA followed by Dunnett’s multiple comparisons test comparing groups to the PDMS_200kPa control condition. e, Principal component analysis of quantified FBR components (αSMA, collagen I, CD68, and capsule thickness). Samples of the same implant stiffness are grouped within a colour-coded envelope. All implants at or below the stiffness of subcutaneous muscle tissue (PAA_20kPa, PDMS_2kPa, PAA_0.2kPa) grouped together away from the highest stiffness PDMS_200kPa. This clustering occurred in the direction of the Dex group, indicating that all three soft coatings considerably alleviate FBR.

The coatings consisted of either soft 0.2 kPa polyacrylamide (PAA_0.2kPa), 2kPa silicone (PDMS_2kPa), or 20 kPa polyacrylamide (PAA_20kPa). One group of implants remained non-coated (PDMS_200kPa) (Supplementary Fig. 4). We also benchmarked the performance of our soft-coated implants against currently used clinical strategies, exploiting devices impregnated with chemical repressors of inflammatory reactions such as glucocorticoids^15, 34^, by including a group of implants coated with a ∼100 µm thick dexamethasone-impregnated silicone of ∼ 200 kPa (Dex)^15^.

These devices were implanted first into the subcutaneous space of rats, a common location for medical implants such as pulse generators^35^ and biosensors^17^, facing the underlying striated muscle. Three months post-implantation, a time point by which acute inflammation due to implantation has resolved and chronic responses to implanted materials have settled in, FBR was significantly reduced around the implants with soft coatings if compared to the stiffer materials as revealed by immunohistochemistry (Fig. 3b). The intensities of markers for myofibroblasts (αSMA), collagen I, and macrophages (CD68) were significantly lower in tissues surrounding soft materials (αSMA *p =* 0.005, collagen *p* = 6.0e-4, CD68 *p* = 3.4e-5, one-way ANOVA) (Fig. 3c). The reduction in FBR around soft implant surfaces was similar to that observed in dexamethasone-treated implants (*p* > 0.05 for all three stains, Bonferroni-corrected Student’s t-tests). Also capsule thickness, a measure of fibroblast proliferation around the implant and of the severity of FBR, was significantly decreased around implants with soft coatings of *G* ≤ 20kPa (Fig. 3d, Supplementary Fig. 6) (*p* = 0.006, one-way ANOVA). Like our *in vitro* results, the extracellular matrix protein fibronectin did not vary significantly across groups (Supplementary Fig. 5) (*p* = 0.14, one-way ANOVA).

Principal component analysis of the analysed FBR markers showed that all three soft coatings clustered together and were distinct from the stiffest coating (Fig. 3e), suggesting that coatings with a shear modulus of up to ∼20 kPa, which is similar to muscle tissue stiffness^31, 36^, ameliorated FBR to subcutaneous implants. These results indicated that FBR can be minimized by coating implant surfaces with a material whose stiffness does not exceed that of the surrounding tissue.

To test if the suppression of FBR by implants with soft coatings is a tissue type-specific phenomenon or if it is more generally applicable, we then implanted nerve conduits with or without soft coatings as those described above in a rat model of nerve injury (Fig. 4a) (for details on sciatic nerve transection see Methods). Similar to the subcutaneous implants, intensity of markers for myofibroblasts (αSMA) and activated macrophages (CD68) were significantly decreased in tissues exposed to a soft coating compared to the non-coated control implants (Fig. 4b, c) (αSMA *p =* 0.002, CD68 *p* = 0.007, one-way ANOVAs). Collagen I also showed a similar trend, although differences were not statistically significant (Fig. 4b, c) (*p* = 0.19, one-way ANOVA), while capsules became significantly thinner around softer materials (Fig. 4d) (*p* = 0.0008, one-way ANOVA). The decrease in the cellular components of FBR around soft implant surfaces was also similar to that of dexamethasone-treated implants (for all soft-coated implants compared to Dex: *p* > 0.05, Bonferroni-corrected Student’s t-tests, Fig. 4b, c), indicating that soft coatings of implants may indeed represent a general approach to alleviate FBR to biomedical implants irrespective of the type of host tissue.

**Fig 4.**
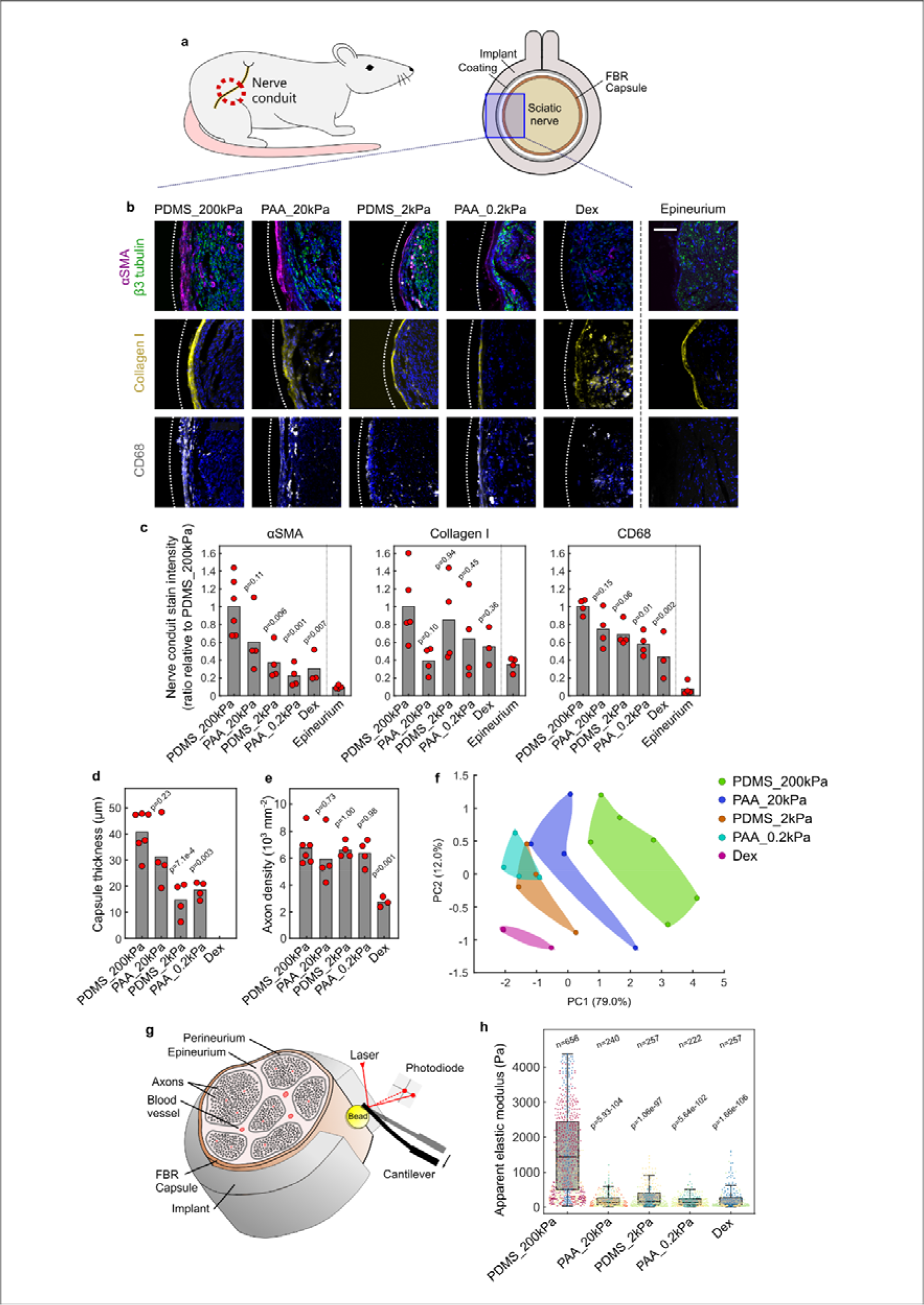
Coatings at least as soft as host tissue significantly reduce foreign body reaction to nerve conduits 3 months post-implantation *in vivo*. a, Schematic of implant location and structure. Implants had an internal diameter of 1.5 mm. Blue square represents area shown in images in b. b, *Z*-stack confocal images of sciatic nerve tissue regenerated through conduits of 200 kPa shear modulus silicone PDMS (PDMS_200kPa), or conduits coated with 20 kPa polyacrylamide hydrogel (PAA_20kPa), 2 kPa PDMS (PDMS_2kPa), 0.2kPa polyacrylamide (PAA_0.2kPa), or PDMS impregnated with 10 mg/ml dexamethasone (Dex). Approximate edges of implants are indicated by dotted white lines. Tissue is fluorescently labelled for axons (β3 tubulin, green), myofibroblasts (αSMA, magenta), extracellular matrix components (collagen I, yellow), and macrophages (CD68, white). All images are stained for nuclei (DAPI, blue). Images of non-operated naïve nerves are also included for comparison (epineurium). Softer coatings reduced cell activation and fibrotic capsule thickness while permitting nerve regeneration. Dex-treatment, however, abolished not only the fibrotic capsule but also nerve regeneration. Scale bar: 100 μm. c, Plot of relative stain intensities. e, Plot of axon density in nerves 5 mm downstream of implantation site. d, Plot of fibrotic capsule thickness. Quantification of thickness in Dex group absent as dexamethasone impeded formation of a structured boundary between tissue and implant. For all plots: bars represent mean, dots represent individual animals. *N* = 3 to 6 rats. f, Principal component analysis of quantified FBR components (αSMA, collagen I, CD68, and capsule thickness). Samples of the same stiffness condition are grouped within a colour-coded envelope. The implants of or below nerve tissue stiffness (PDMS_2kPa, PAA_0.2kPa) grouped together, away from the higher stiffnesses PAA_20kPa and PDMS_200kPa, and in the direction of the anti-inflammatory Dex group, indicating that the softest PDMS_2kPa and PAA_0.2kPa most efficiently alleviated FBR g, Diagram of *ex vivo* AFM setup for nerve tissue stiffness measurements. h, Box and scatter plot of tissue stiffness values for tissue in proximity to the implants with various coatings, showing a significant stiffening of tissue around implants with a stiff surface. Tissue stiffness was similar around PDMS_2kPa, PAA_0.2kPa, and Dex-treated implants (*p* > 0.05). *n* = number of measurements. Measurements taken on *N* = 3 to 6 rat nerves; measurements taken from the same animal are depicted with the same colour. All statistical comparisons carried out via one-way ANOVA followed by Dunnett’s multiple comparisons test comparing groups to the PDMS_200kPa control condition. Bonferroni-corrected Student’s t-test used for comparisons to Dex group. Box plot characteristics described in Methods.

Principal component analysis of the analysed FBR markers revealed that, as in the subcutaneous model, the softer implants were closer to the implants treated with dexamethasone, suggesting that the softest coatings ameliorated FBR most. In contrast to the subcutaneous model, however, in nerves only the two softest coatings (*G* = 0.2 kPa and 2 kPa) clustered together and were distinct from the stiffer materials of *G* = 20 kPa and 200 kPa (Fig. 4f). Because nerves are softer than subcutaneous tissue, with a stiffness on the order of ∼1 kPa^24^, our results confirmed that FBR may indeed be minimized by implant coatings at least as soft as the surrounding host tissue.

While soft-coated implants exhibited similar anti-FBR properties as dexamethasone-treated materials, glucocorticoids such as dexamethasone are not only anti-inflammatory but also anti-proliferative. Consequently, neuronal regeneration was significantly reduced in dexamethasone-doped nerve implants if compared to implants with soft coatings (Fig. 4b, e, Supplementary Fig. 7) (*p* = 0.003, one-way ANOVA). Hence, our data showed that soft coatings of implants not only suppress inflammation and FBR but also permit regeneration – in contrast to currently exploited devices relying on the anti-inflammatory properties of glucocorticoids.

In FBR and fibrosis, the secretion of dense networks of extracellular matrix rich in components such as collagen I leads to an increase in the stiffness of the tissue surrounding the implant^37, 38^. To further corroborate the suppression of FBR in tissues surrounding implants with soft coatings, we used *ex vivo* atomic force microscopy to measure the apparent elastic moduli of tissues exposed to the different implants (Fig. 4f). With a median reduced apparent elastic modulus *K* ∼ 1200 Pa, tissue around the stiffest conduits showed a significant degree of stiffening compared to tissue around all soft-coated and dexamethasone-treated implants with *K ∼* 150 Pa (p < 0.001 relative to all soft-coated and dex-treated groups, Dunnett’s multiple comparison test), which were indistinguishable from each other (*p* = 0.70; Bonferroni-adjusted Student’s t-test) (Fig 4g), adding further evidence that soft implant coatings minimised the development of FBR.

Both our *in vivo* and *in vitro* experiments indicated that different implant coatings considerably impacted the onset and extent of FBR. However, the different materials used in this study might not only differ in their mechanical properties but also in their ability to bind proteins. Protein adsorption – an initial and critical step in FBR – can vary between materials and lead to differences in FBR severity^39^. Hence, we next tested if protein adsorption depended on the type of material using a bicinchoninic acid (BCA) assay. We found no differences in protein adsorption across our different surfaces (Supplementary Fig. 8) (*p* = 0.954, one-way ANOVA), suggesting that FBR to implants stiffer than the host tissue was indeed initiated by a direct, mechanosensitive response of cells rather than indirectly via alterations in protein adsorption.

To further corroborate that cellular mechanotransduction – the conversion of mechanical cues into biochemical signals – was involved in FBR to implants stiffer than the host tissue, we investigated the cellular distribution of the transcriptional regulator YAP in tissue surrounding implants after 3 months. In many systems, YAP is majorly involved in mechanotransduction^40^. On soft substrates, YAP is usually excluded from the nucleus, while on stiffer substrates YAP enters the nucleus, leading to mechanically driven changes in gene expression^41^. This effect has been observed *in vitro* in several cell types including fibroblasts^42^ as well as *in vivo*^43^. Immunohistochemical stains indeed revealed a significantly lower nuclear localisation of YAP in tissues exposed to softer coatings if compared to stiffer coatings in both implant types (Fig. 5) (subcutaneous *p* = 0.01, nerve *p* = 0.001, one-way ANOVAs), confirming that cells in the vicinity of medical implants sense and respond to the stiffness of the implant material.

**Fig 5.**
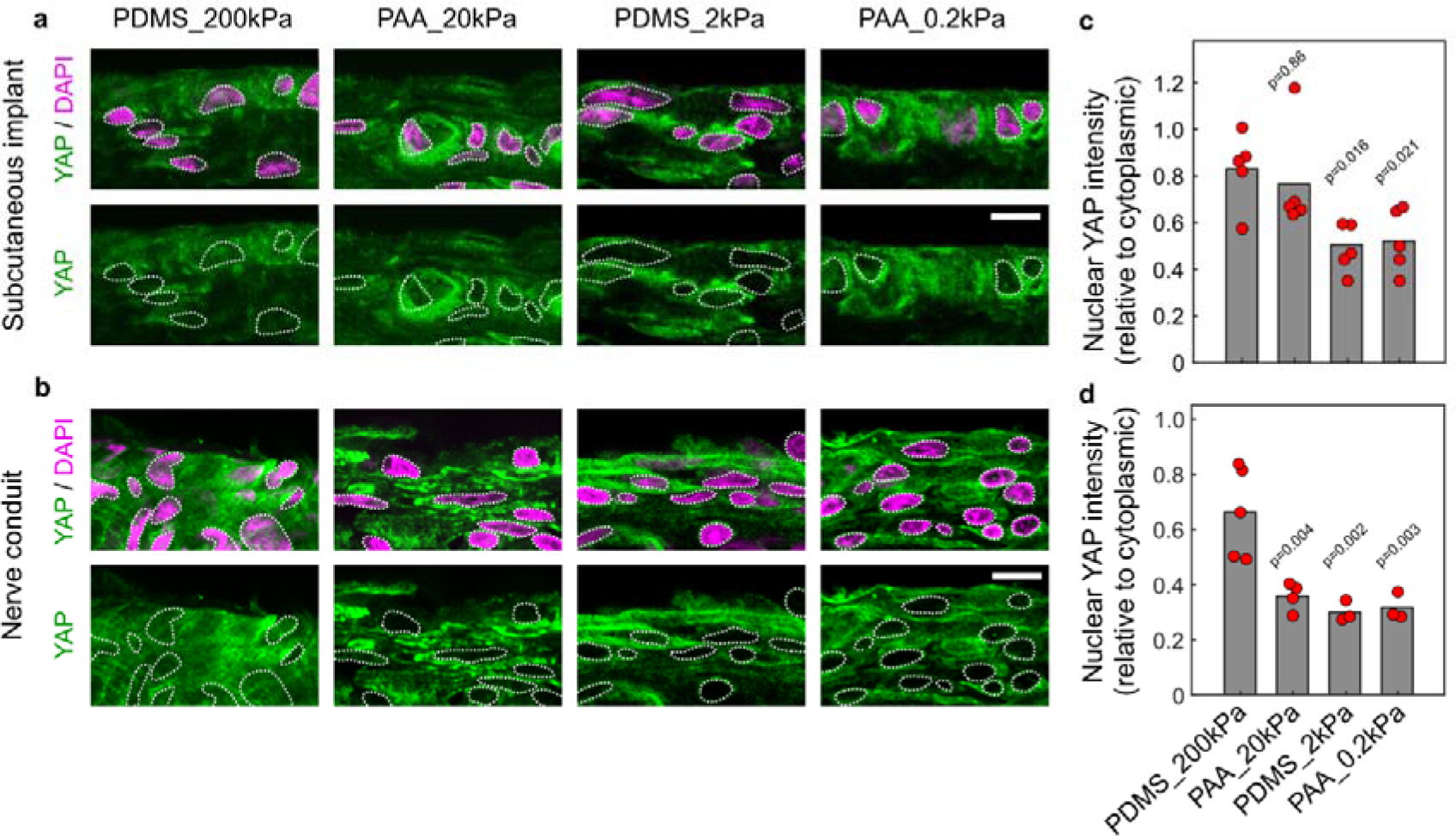
Reduction in foreign body reaction *in vivo* correlates with exclusion of the mechanosensitive transcriptional regulator YAP from nuclei in tissues surrounding soft-coated implants. a-b, Confocal images of subcutaneous (a) and nerve tissue (b) stained for the transcriptional regulator YAP and nuclei (DAPI). Tissue was exposed to implants of 200 kPa shear modulus silicone PDMS (PDMS_200kPa), or to implants coated with 20 kPa polyacrylamide hydrogel (PAA_20kPa), 2 kPa PDMS (PDMS_2kPa), or 0.2 kPa polyacrylamide (PAA_0.2kPa), for 3 months. YAP preferentially localised into nuclei of cells in contact with stiff but not with soft-coated implants, indicating a mechanosensitive process. Nuclei delineated by dashed white lines. Scale bars: 10 μm. c,d, Plots of relative nuclear YAP intensities. Bars represent mean, dots represent individual animals. *N* = 5 rats (c) and *N* = 3 to 5 rats (d). All statistical comparisons carried out via one-way ANOVA followed by Dunnett’s multiple comparisons test comparing groups to the PDMS_200kPa condition.

## Discussion

As in previous *in vitro* studies of similar cell systems, primary bone marrow-derived macrophages and peripheral nerve tissue-derived fibroblasts investigated in this study responded to the stiffness of their environment^17, 44–47^. Our results link this cellular mechanosensitivity to a key problem in medical implants: FBR. Contact with materials stiffer than native tissue led to trans-differentiation of fibroblasts into myofibroblasts, and macrophages became activated and took on an M2-like phenotype, driving tissue generation an thus leading to thicker fibrotic capsules. This effect was similar for two different soft materials: polyacrylamide and PDMS, and it occurred irrespective of implant design and the location of implantation, indicating a very robust response of host tissues to implant stiffness.

Collectively, our results show that a mechanical mismatch between implant material and host tissue stiffness may drive FBR at the cellular level. In nerves – a tissue with a stiffness on the order of ∼1kPa^24^ – coatings of 2kPa and 0.2 kPa were most effective in minimising FBR (Fig. 4f), while in the subcutaneous space – a tissue with a stiffness of approximately 12 kPa^31, 36^ –all coatings not exceeding 20 kPa were similarly effective (Fig 3e). Our data are consistent with recent observations that ultra-soft implants can reduce FBR compared to stiffer counterparts^17, 19^, and they define the upper limit of implant stiffness softening an: implant surface to at least match the stiffness of its host tissue resulted in maximum reduction of mechanically triggered FBR. Additionally, by using implants with similar structural mechanical properties and various thin soft coatings, we showed that this threshold of FBR is determined at the cellular level and not a result of the mechanical mismatch between tissue and the whole implant as generally thought^20–22^.

Mechanotransduction – the process by which cells translate mechanical cues into biochemical responses – plays a key role in many cellular processes^48^. Cells constantly probe their mechanical environment by applying contractile forces. This activates downstream signalling pathways – which have yet to be fully characterised – leading to changes in gene expression and cell function. Mechanotransduction has recently been associated with inflammation^17^ and fibrosis^19^ *in vitro*. Here, we provide direct evidence for an involvement of cellular mechanotransduction in the regulation of immune cell and connective tissue cell function *in vivo*.

The transcription factor yes-associated protein (YAP) is a key downstream mediator in many mechanotransduction pathways^40, 41^. Its substrate stiffness-dependent translocation into the nucleus is a useful marker to identify systems in which mechanotransduction is playing an active a role^40–43^. We observed that, in tissue surrounding an implant, YAP’s cellular distribution is affected by implant stiffness *in vivo*. While softer materials have been used in the past to decrease tissue damage around implants and to increase their biocompatibility^20, 49–51^, our results provide a first direct link between *in vivo* cellular mechanotransduction and FBR.

Current medical implants offer a powerful tool for the treatment of a number of clinical conditions. However, long term stability of implants often remains limited by FBR, particularly of those actively interacting with the host’s environment such as electrical neural interfaces. Corticosteroid drugs such as dexamethasone offer a viable strategy to control FBR in certain situations^34^. However, their anti-inflammatory effect also greatly interferes with regeneration of surrounding tissue^15^, making them ill-suited for use in regenerative implants. As shown here, coatings at least as soft as the host tissue offer an effective strategy to minimise FBR without impacting surrounding tissue function or sacrificing the bulk mechanical properties of the implant itself.

Medical implants are usually prepared from materials compatible with miniaturisation, surgical handling, and advanced 3D design. These materials are much stiffer than their biological host tissues. Tissue-level soft coatings offer a number of advantages as FBR- minimising components of medical implants. The coatings’ dimensions and compatibility with existing microfabrication techniques make this technology potentially easy to apply to many implant designs, which may facilitate translation to the clinic. Moreover, the FBR-reducing effects are linked to the mechanical – and not chemical – properties of the coating materials, providing great flexibility in material choice for different implant designs and applications. Hence, the optimization of the mechanical surface properties of medical implants, in addition to that of their chemical and electrical properties, will likely greatly contribute to long-term stability of these devices in the future.

## Materials and Methods

### Polyacrylamide cell culture substrates

Polyacrylamide hydrogels were prepared two days prior to the plating of any cells, using a protocol previously described^52^. 19 mm diameter glass coverslips were cleaned by alternate dipping in ddH_2_O and EtOH, and covered in 0.1N NaOH (VWR, E584) for 5 min. NaOH was removed, and coverslips were functionalised with 150 μl of APTMS (3-Aminopropyltrimethoxysilane; Sigma, 281778) solution for 2.5 min, followed by thorough rinsing in water. Coverslips were finally allowed to sit in a glutaraldehyde 0.5% (v/v) solution (G6257, Sigma) in ddH_2_O (15230089, Fisher Scientific) for 30 min at RT.

Polyacrylamide premixes were prepared by mixing of acrylamide (40\% w/w; A4058, Sigma), bis-acrylamide (2%; BP1404-250; Fisher Scientific), and hydroxy-acrylamide (97%; 697931; Sigma) solutions (177:100:23 ratio). A volume of PBS (D8537, Sigma) was added to the premix to achieve gels of a particular stiffness (Supplementary Table 1). This final gel mix was degassed under a vacuum for 10 min.

To initiate polymerisation, 5 μl of ammonium persulfate solution (0.1 g/ml in ddH_2_O; Sigma, 215589) and 1.5 μl of TEMED (N,N,Ń,Ń-tetramethylethylenediamine; 15524-010, Invitrogen) were added to 500 μl of gel mixes. 8 μl drops of mix were placed on the treated surface of 19 mm diameter coverslips, and covered with a 22 mm diameter glass coverslip (previously treated with a RainX hydrophobic coating – Rain Repellent Krako Car Care International Ltd, UK).

The gels were allowed to swell in PBS overnight, allowing any remaining monomers and unreacted crosslinkers to leach out. 22 mm diameter coverslips were then removed to reveal the hydroxyacrylamide gels bound to 19 mm coverslips. Some of the gels were examined at this point under AFM to ensure their stiffness matched the expected values. The rest of the gels were sterilised under UV light for 1 hour, and functionalised with PDL (Poly-D-lysine; 100 μg/ml in PBS; P6407, Sigma) at room temperature overnight. Prior to cell plating, gels were placed in culture medium for 30 min to allow medium to fill them, and prevent cell culture medium from being diluted when cells were added.

Polyacrylamide substrates were prepared in the range of 0.1 kPa to 50 kPa. This covered the range of tissue stiffnesses typically found in the body (with 50 kPa being stiffer than most soft tissues and organs^25^). This was also the range of stiffnesses we could reliably achieve using the same polymer : cross-linker ratio (Supplementary Table 1).

#### In vitro assay

##### Nerve fibroblasts

Cultures were prepared from postnatal day 1 to 5 Sprague Dawley rat (Charles River UK) sciatic nerves. Using a variation of the procedure previously described^53^. All animal procedures carried out were in compliance with the United Kingdom Animals (Scientific Procedure) Act of 1986 and institutional guidelines.

Sciatic nerves of 10 - 20 animals were dissected out using sterilised microscissors and fine forceps, and were kept in chilled HBSS (14170-112, Invitrogen) to stabilise the pH and osmotic environment. To dissociate the tissue, nerves were transferred to a 2 ml collagenase solution (2 mg/ml; C9407, Sigma) and incubated for 30 min at 37 °C, after which 2 ml of trypsin (1 mg/ml; T0303, Sigma) was added (20 min, 37 °C incubation). Finally, 2 ml of deoxyribonuclease (0.1 mg/ml; D5025, Sigma) was added and, after a brief incubation period (2 min), the cells were centrifuged (4 min, 1000 rpm). The supernatant was removed, and the cell pellet was re-suspended in 2 ml of triturating solution (containing 10 mg/ml bovine serum albumin [A7906, Sigma], 0.5 mg/ml trypsin inhibitor [10109886001, Roche], 0.02 mg/ml deoxyribonuclease).

To isolate the population of nerve fibroblasts from the dissociated nerves, the cells were centrifuged and re-suspended in 0.5 ml of DPBS/BSA (Dulbecco’s phosphate-buffered saline [14190-094, Invitrogen] supplemented with 5 mg/ml bovine serum albumin) and 50 μl of 50nm-magnetic-bead antibodies against rat/mouse CD90.1 (Thy1.1) (Miltenyi Biotec, 120-094-523) and incubated for 15 min at RT. While nerve fibroblasts are less well studied than other cell types, they are known to express CD90.1, while other common nerve cell types do not^54^. This made CD90.1 a good choice for selectively isolating this cell population. Cells were centrifuged, re-suspended in 2 ml of chilled DPBS/BSA, and run through a column equipped with a magnetic separator (MiniMACS Separator; Miltenyi Biotec, 130-042-102). Once the buffer had finished running through the column, and the flow-through collected, the column was removed from the magnetic separator, and the magnetically-labelled cells were flushed out with chilled DPBS/BSA. Finally, the positive fraction (nerve fibroblasts) were centrifuged and the cell pellet re-suspended in DMEM (11320-033, Invitrogen) supplemented with a further 4 mM of glutamine (25030032, Invitrogen), 100 mg/ml foetal calf serum (FCS, Invitrogen) and an antibiotic-antimycotic agent (15240-062, Invitrogen). The cells were plated on polyacrylamide substrates at a density of 10,000 cells/cm^2^. While these 50nm beads remain adhered to the cells of the positive fraction, they are not known to lead to lead to changes in cell behaviour.

##### Bone marrow-derived macrophages

Macrophages were derived from adult rat bone marrow hematopoietic stem cells as previously described^55^. Adult Sprague Dawley rats (Charles River UK) were sacrificed by exposure to a rising concentration of CO_2_. The femurs were dissected out and broken open using sterile scissors. Bone marrow contained within the femurs was washed out with chilled DPBS (14190-094, Invitrogen) and collected. The cell suspension was centrifuged for 10 min at 1000 rpm. The supernatant was discarded and the cells were re-suspended in BMDM (the same supplemented DMEM used for nerve fibroblasts, further supplemented with macrophage colony stimulating factor [400-28, Peprotech; 50 ng/ml]). Cells were counted in a hemocytometer and seeded on 100 x 15 mm uncoated petri dishes (Sigma, P5731) at a density of 5,000 cells/cm^2^. The dishes were supplemented with further medium after 3 days of culture at 37 °C.

Hematopoietic stem cells differentiate into macrophages in the presence of macrophage colony stimulating factor present in BMDM. After 6 days to allow differentiation of Hematopoietic stem cells differentiate into macrophages, the medium in the dishes was removed to dispose of any cells which had not attached to the substrate. The remaining cells were washed with warm DPBS followed by CellStripper solution (Corning, 25-056-CI). Cells were incubated in CellStripper for 5 min at 37 °C to detach them from the substrate. The cell suspension was collected and an equal volume of BMDM was added to inactivate the CellStripper solution. Cells were then centrifuged at 1000 rpm for 10 min and the supernatant was removed. Upon re-suspension in BMDM and counting, cells were plated onto polyacrylamide substrates at a density of 10,000 cells/cm^2^.

### Immunocytochemistry

Cell stains were carried out 6 days after plating on polyacrylamide substrates. At this point in time, warm paraformaldehyde solution (40 mg/ml in PBS) was added to the cells for 15 min at room temperature. The fixative solution was washed off with PBS (3 washes, 10 min per wash). To improve antibody specificity cells were incubated for 30 min at room temperature in a blocking solution consisting of 0.03% v/v Triton X-100 (Sigma, T8787) and 3% v/v bovine serum albumin (Sigma, A9418) in PBS. Primary antibodies (in blocking solution) were then added to cells and incubated overnight at 4 °C. Further details regarding antibody concentrations can be found in Supplementary Table 2.

Excess primary antibodies were washed off using PBS (3 washes, 10 min). Secondary antibodies in blocking solution were incubated on the cells for 2 hr at room temperature. Following two washes with PBS and a wash with non-saline Tris-buffered solution, Fluorsave mounting agent (Millipore, 345789) was added to sections to preserve fluorescence before gels were placed onto glass slides and stored at 4 °C prior to imaging.

Imaging of stained cells on polyacrylamide substrates was carried out using a confocal microscope (Leica TCS SP5). For every condition 3 images were taken at random sites within each gel with a 20X objective. The nuclear stain was used as a guide to ensure cells were present in the field of view when pictures were taken. Gain and exposure settings for each channel used were maintained constant between imaging sessions and across different stains.

Cell counts were performed by hand in the Image-J software package (v1.48, National Institutes of Health, USA). Contrast of images was modified prior to analysis. For morphological stains, one image of high stiffness (50 kPa) and one of low stiffness (0.1 kPa) were opened and their contrast modified to an equal degree until a satisfactory pattern of stain was achieved in both. This contrast modification was then applied to all images of the same batch of stained gels. For cell-type specific stains such as alpha-smooth muscle actin this same contrast modification was carried out using negative and positive control stains. Cells were counted, or the area stained was designated, by hand. Statistical analysis and data plotting was carried out using MATLAB (Mathworks, R2016b). Box plots used to display data consist of a box containing the 25^th^ to 75^th^ percentile range of data (the interquartile range), an inner box mark indicating the median value, and whiskers containing most of the remaining data points. Outliers, defined as data points further away from the 25^th^ or 75^th^ percentile than 1.5 times the interquartile range, are presented as circles. Box plots with overlaid scatter plots do not depict these outliers, as these can be observed in the scatter plot.

### RNA sequencing

RNA sequencing was performed in parallel on n = 4 biological replicates. RNA extraction was carried out at day 3 of culture on polyacrylamide substrates using an RNeasy Plus Micro Kit (Qiagen, 74034). This time point (different to the 6 day time point used in immunocytochemistry experiments) was chosen as it was the point at which cells were beginning to show major morphological differences across conditions, while minimising the contribution of potential changes in gene expression due to other effects - such as increased proliferation rates and subsequent increases in cell density seen on stiff substrates. Prior to and at regular intervals during the extraction procedure, work surfaces and pipettes were cleaned with RNase Zap decontamination solution (ThermoFisher, AM9780) to inactivate RNases and prevent sample RNA degradation. To collect the cells, each coverslip was briefly washed in PBS and lifted out of the solution using forceps. The gels onto which the cells were attached were gently scraped off from the coverslips using a sterile steel blade and placed in RLT lysis buffer plus (Qiagen). The samples in buffer were then moved to QIAshredder tubes (Qiagen, 79654) and centrifuged in a microcentrifuge (MSE, mistral 1000) for 2 min at 8,000g. Instructions provided by the kit manufacturer were then followed to extract cellular RNA, which was collected and stored at -80°C.

RNA quantification and integrity analysis were carried out on all samples prior to library preparation. Using an RNA 6000 Pico Kit (Agilent, 5067-1513), samples concentration and integrity was analysed using an Agilent 2100 Bioanalyzer. No samples with an RNA integrity number <7 were used. Library preparation was thereafter carried out using an Ovation RNA-Seq System V2 kit (NuGen, 7102-32), following manufacturer instructions. Samples were finally submitted for sequencing in an Illumina HiSeq 2500 system.

Illumina read data files were run through a bioinformatics pipeline and aligned with the *Rattus norvegicus* genome (Ensembl Rnor_6.0). Fold changes and p-values were calculated for each gene between each experimental group and a control group (consisting of the combined 0.1 kPa and 1 kPa conditions). Genes expressed were filtered to produce lists of differentially expressed genes (DEGs). Defined by a minimum of 2-fold change in expression, a base expression above 3 normalised counts, and an adjusted p-value below 0.05. RNAseq data have been deposited in the ArrayExpress database at EMBL-EBI (https://www.ebi.ac.uk/arrayexpress/experiments/E-MTAB-7900) under accession number E- MTAB-7900. Code to create the figures displaying RNAseq results is available in the following GitHub repository: https://github.com/CTR-BFX/2019-Carnicer-Lombarte.

### Implant fabrication

#### Nerve conduits

Moulds of the conduit implants were designed in 3D-CAD software (AutoCAD, Autodesk Inc) and 3D printed in PLA (polylactic acid) plastic using a MakerGear M2 3D printer (MakerGear). The moulds were covered in Sylgard 184 PDMS (Dow Corning), and were placed in an oven at 65°C overnight. The 3D printed mould was removed from the cured PDMS, producing a negative pattern of the conduit implants. The surface of the PDMS negative moulds was activated using oxygen plasma (Diener plasma etcher) for 25 seconds at 30 W, and 0.8 mbar chamber pressure. PDMS moulds were then functionalised using Trichloro(1H,1H,2H,2H-perfluorooctyl)silane (Sigma, 448931). A few drops of silane were placed on a glass petri dish and into a desiccator together with the PDMS moulds. The desiccator was pumped down into a vacuum, and functionalisation was allowed to take place overnight. The resulting layer of silane prevented any new PDMS cured on these moulds from binding to them, allowing for the casting of the PDMS conduits from these negative moulds.

To cast the conduits, flat petri dishes with raised edges were prepared. These edges were produced through layering multiple layers of insulation PVC tape, until a thickness of 0.6 mm was achieved. The functionalised PDMS negative moulds were coated with a thick layer of Sylgard 184 PDMS and placed on top of these dishes. The raised edges of the dishes created a 0.6 mm thick layer of Sylgard 184 PDMS below the moulds, which would become part of the conduits after curing. The moulds and dishes were placed in an oven at 65°C overnight.

The freshly-cured layer of PDMS was carefully peeled from the moulds and trimmed to the appropriate dimensions with a steel blade. The resulting PDMS implants were functionalised with an additional layer of silicone/polyacrylamide before rolling into conduits. To roll into conduits, the edges of the implants were brought together and carefully secured with insulation PVC tape. The edges were covered with RTV PDMS (SA03073, Farnell), which was allowed to cure overnight. An additional layer of RTV was then added and allowed to cure before the conduits were stored in PBS and sterilised under UV prior to implantation. The resulting conduit had a length of 7 mm, an internal diameter of 1.5 mm, and a wall thickness of 0.6 mm. Supplementary Fig. 9 summarises the conduit fabrication protocol.

#### Subcutaneous implants

A 3 mm thick layer of Sylgard 184 PDMS was cast and cured overnight at 65 °C. This was then trimmed using a steel blade into 5 x 5 mm blocks. Each block then was functionalised with a coating. Four blocks – one for each of the 4 stiffness-controlled conditions – were combined into one 10 x 10 mm implant and stuck together using RTV silicone. The sides of each of the four component blocks were notched to later be able to identify them. Dexamethasone-doped implants remained as 5 x 5 mm blocks, and were not combined with other implants. Implants were stored in PBS and sterilised under UV prior to implantation.

#### Coatings

To produce dexamethasone-doped silicone implants (Dex), Sylgard 184 PDMS was doped with 10 mg/ml of dexamethasone and spin-coated into 100 μm-thick films. The dexamethasone-doped films were cut into appropriately-sized rectangles. A small amount of RTV PDMS was spread over an implant and a dexamethasone-doped PDMS rectangle was placed on top. After allowing the RTV to cure overnight, the dexamethasone-doped film was further trimmed to match the shape of the underlying PDMS, and the PDMS conduitwas rolled as described above.

Soft silicone coatings (PDMS_2kPa) were prepared from a mix of NuSil 8100 (Polymer System Technology, GEL-8100) and Sylgard 184 (99% to 1% w/w, respectively). An implant was thoroughly cleaned with ethanol and ddH_2_O (15230089, Fisher Scientific) and dried with nitrogen gas, followed by the application of a 9 μl drop of the soft silicone mix to its surface. This drop was spread out to ensure that the entire inner surface of the implant was completely covered. The implant was then transferred to an oven and baked at 65 °C for one week. This long curing time was a necessary step to remove traces of non-cured PDMS.

Polyacrylamide coatings (PAA_0.2kPa and PAA_20kPa) were grafted onto Sylgard 184 PDMS implants following a published protocol^56^.Glass coverslips were cleaned by alternate dipping in ddH_2_O and EtOH. PDMS implants were thoroughly cleaned with methanol, dried with nitrogen gas, and covered with a benzophenone solution 10% w/w in EtOH (Sigma, B9300) for 2 min at room temperature. Benzophenone was removed and conduits/subcutaneous implants cleaned with methanol and dried with nitrogen gas. Polyacrylamide hydrogel mixes were prepared by combining acrylamide and bisacrylamide solutions at a 2:1 ratio. The mixes were combined with PBS to achieve the desired stiffness, as described in Supplementary Table 1. To initiate the polymerisation of the gel mix, 5 μl of APS solution (0.1 g/ml in ddH_2_O) and 1.5 μl of TEMED were added to 500 μl of gel mixes. A 9 μl drop of the gel mix was then transferred to a PDMS implant, which had been previously soaked in 10% (w/w) benzophenone solution in ethanol. The drops were spread out by covering with a clean glass coverslip. Implants were then quickly transferred under a 12 J/cm^2^ UV lamp for 10 min (SUSS MicroTec MJB4). After the 10 min of UV exposure, glass coverslips were removed and polyacrylamide-PDMS composites were placed in PBS for a further 30 min. The polyacrylamide coating was kept hydrated with PBS at all times until implantation.

Stiff silicone implants (PDMS_200kPa) were not coated with anything, leaving the surface of the Sylgard 184 implant exposed to the tissue. While the absence of a coating resulted in slightly larger diameter conduits for this group, this was unlikely to have an impact on the nerve tissue. Conduits were implanted into a nerve gap injury which was later filled with nerve tissue as part of normal tissue regeneration (and not wrapped around a pre-existing nerve).

### *In vivo* implantation

All experimental procedures were performed in accordance with the UK Animals (Scientific Procedures) Act 1986. Surgical procedures were carried out under aseptic conditions. ∼250 g Lewis rats (Charles River UK) were housed in groups of 5 and provided *ad libitum* access to food and water for a minimum of 7 days prior to surgical procedures. Immediately prior to all surgical procedures, animals received an injectable dose of the non-steroidal anti- inflammatory drug meloxicam (1.5 mg/ml, subcutaneous). Anaesthesia was induced and maintained with isoflurane delivered via a facemask. Body temperature was monitored via a rectal probe and maintained at 37 °C using a thermal blanket.

#### Nerve conduits

Biceps femoris and vastus lateralis muscles of the right leg of the animals were approached dorsally and separated to expose the septum through which the sciatic nerve travels. The sciatic nerve trifurcation point was located and followed 2 mm proximal. This site was used as a landmark to achieve consistent location of injury and implantation. The nerve was cleanly transected at this location using scissors and the conduit was positioned between the two resulting nerve stumps, leaving a 5 mm long empty gap within the conduit between the stumps. The epineurium of each nerve stump was sutured to the silicone tube using 9/0 nylon sutures (Ethicon). Each animal received only one conduit, with a single type of coating.

#### Subcutaneous implants

An incision was done dorsally over the right leg of an animal (approximately above the femur). The skin was separated from the underlying muscle fascia using blunt forceps to create a tunnel from the site of incision towards the midline of the animal. The implant was fed through this tunnel and placed ∼1 cm away from the midline, with the coating facing the muscle fascia. Each animal received one subcutaneous implant; either a composite of all four stiffnesses or a dexamethasone-containing implant. This was done to prevent any dexamethasone leached from the implant from interfering with the FBR of all other conditions.

All animals were allowed to recover following implantation. A further dose of meloxicam was given orally the day after surgery. 3 months post-implantation, animals were sacrificed by exposure to a rising concentration of CO_2_ or, if AFM measurements were to be carried out, by anaesthetic overdose (see below)and the tissue collected.

### Immunohistochemistry

All tissue was fixed prior to processing and staining by immersion in paraformaldehyde solution (40 mg/ml in PBS) overnight at 4 °C. Samples which required sectioning were then transferred to a sucrose solution (30% w/w in PBS; S0389, Sigma) for cryoprotection. They were kept in this solution for a minimum of 16 hr at 4 °C, and otherwise stored until further processing. Cryopreserved samples were embedded in optimal cutting temperature compound (Tissue-Tek, 4583), which was frozen and mounted on a cryostat (CM3050 S, Leica). 12 μm - thick sections were cut from the samples at a cutting temperature of -20 °C. Sections were placed on glass slides and allowed to dry at room temperature overnight before storage at -20 °C until stained.

Sections ready to be stained were washed in a Triton X-100 0.1% v/v solution in PBS to allow for permeabilisation. These and all further washes were performed three times for 10 min. To minimise non-specific antibody binding, sections were incubated in a blocking buffer, consisting of tris-buffered saline containing 0.03% v/v Triton X-100 and 10% v/v donkey serum (Millipore, s30-100ml). After blocking for 1 hr at room temperature, primary antibodies (in 10% blocking buffer v/v in PBS) were added to sections (further details in Supplementary Table 2). Sections were covered with paraffin film to prevent drying and were incubated in primary antibodies overnight at 4 °C.

Sections were washed in PBS-Triton solution to remove excess primary antibodies, and then incubated in secondary antibodies in the same solution as primaries for 2 hr at room temperature. Secondary antibodies were finally washed off with two PBS washes followed by a non-saline Tris-buffered solution (T6066, Sigma). Fluorsave mounting agent (Millipore, 345789) was added to sections to preserve fluorescence before encasing with a glass coverslip and storing at 4 °C prior to imaging.

Imaging of stained nerve tissue was carried out using a confocal microscope (Leica TCS SP5). Image files were exported and processed for analysis in Image-J software package (v1.48, National Institutes of Health, USA). Stain intensity profiles of FBR capsules was carried out through a combination of custom Matlab and Fiji scripts. The edge of the nerve capsule was delineated by the user and aligned by the scripts. An intensity profile (intensity vs. depth into the nerve) of the each stain was obtained. The average intensity from the edge of the nerve to a depth of 25 μm was calculated and provided as a ratio to the same intensity of the PDMS_200kPa group. The only exception were CD68 stains, were a depth of 50 μm was instead chosen as macrophages were found to mostly locate deeper into the tissue than other markers. Capsule thickness was analysed using a Matlab script, after its edge was marked by hand based on the αSMA stain. Principal component analysis of FBR was done over the combination of αSMA, collagen I, CD68 stains and capsule thickness. For this analysis capsule thickness in dexamethasone group was set as 0. All values were normalised and missing values substituted for the mean of the group (10 of 100 values in subcutaneous samples, 3 of 82 values in nerves). Axon density was analysed in an automated fashion using a Fiji script over 3 randomly chosen 100 x 100 μm boxes for every image. This was done 5 mm distal to the point of conduit implantation to minimise any effects from the conduits on regeneration, as this could result in a non-uniform axon distribution. Statistical analysis and data plotting was carried out using MATLAB (Mathworks, R2016b).

#### BCA protein adsorption assay

Adsorption of proteins to the surface of materials used in *in vivo* experiments was determined using a BCA assay. Blocks of polyacrylamide hydrogel or silicone rubber were prepared with dimensions 5 x 5 x 2 mm. Blocks were soaked in 10 mg/ml foetal calf serum (FCS, Invitrogen) for 2 hours at 37 °C. The blocks were then removed and placed in 50 mM ammonium bicarbonate solution for a further two hours at room temperature to extract adsorbed proteins. A bicinchoninic acid (BCA) assay (ThermoFisher, 23225) was used to quantify protein content following manufacturer specifications. Absorbance of the assay was measured using a FLUOstar Omega microplate reader (BMG Labtech) at 570 nm, and compared to that of a protein standard.

### Atomic force microscopy

Sample elasticity was determined via atomic force microscopy (AFM) as previously described^57^. Indentation measurements using a cantilever probe were taken on samples placed on an inverted optical microscope (Axio Observer.A1, Carl Zeiss Ltd.) using a JPK Nanowizard Cellhesion 200 AFM (JPK Instruments AG). Tipless silicon cantilevers (Arrow- TL1; NanoSensors) with a spring constant of ∼0.02 N/m were used in experiments were tissue stiffness was measured. Material characterisation made use of either these or stiffer cantilevers (SICON-TL-20, spring constant ∼0.29 N/m, AppNano; TL-FM-10, spring constant ∼2.8 N/m, Nanosensors; spring constant value was measured for each individual cantilever via the thermal noise method^58^). Each cantilever had a polystyrene bead (∼37 μm diameter for tissue, ∼20μm for stiffer materials; Microparticles GmbH) glued (ultraviolet curing, Loctite) prior to all measurements. Choice of cantilever spring constant and bead diameter was taken to ensure high measurement sensitivity for a given indentation depth given the mechanical properties of the sample.

#### Tissue preparation

Lewis rats (Charles River UK) were sacrificed by overdose of euthatal (pentobarbitone) administered intraperitoneally, followed by neck dislocation. Euthatal was combined at a 1:1 v/v ratio with lidocaine anaelgesic and delivered at a total dose of 3 ml/kg of bodyweight. Further processing of tissue was carried out within 1 - 2 hr of animal sacrifice, and all AFM measurements were completed within 5 hr to minimise tissue degradation. This sacrificial method was chosen instead of rising CO_2_ concentration, as changes in tissue stiffness as a consequence of changes in tissue pH can occur with the latter^59^.

To dissect out the sciatic nerve, the dorsal side of the hindlegs was exposed and skin removed. Biceps femoris and vastus lateralis muscles were separated to expose the septum through which the sciatic nerve travels. The nerve was dissected out and transected just below the trifurcation point, and 1 cm above it. The nerve was then transferred to a dish containing mammalian physiological saline previously described by others^60^ (121 mM NaCl, 5 mM KCl, 1 mM MgCl2, 1 mM CaCl2, 0.4 mM NaH2PO4, 23.8 mM NaHCO3, 5.6 mM glucose). Mammalian physiological saline was prepared freshly prior to the experiment. To establish a pH of 7.3, a gas mixture of 95% CO_2_ and 5% O_2_ was bubbled through the solution.

For naïve nerve measurements, under a dissection microscope, blood vessels and excess tissue surrounding the nerve were removed, and the nerve stumps were trimmed off with a steel blade. The remaining nerve was then cut into several fragments for mounting and sectioning. Nerve fragments were embedded in warm 4% w/w low melting point agarose (A9539, Sigma) in PBS. Agarose was allowed to cool and harden for a few minutes before trimming into blocks containing the nerve fragments. Blocks were stuck to a steel stage with cyanoacrylate glue (Super Glue, Loctite) and transferred to a chamber filled with chilled mammalian physiological saline. The blocks were cut into 500 μm thick sections in a vibrating microtome. Nerve sections were transferred to mammalian physiological saline solution containing the live stain fluoromyelin (1:250 v/v in mammalian physiological saline; Invitrogen, F34651) and incubated for 1 hr at room temperature to stain the myelin surrounding axons. This allowed the endoneurial compartment to be identified and later probed via AFM. Sections were mounted onto 35 mm plastic dishes (Z707651, Sigma). The sections were gently deposited onto two strips of cyanoacrylate glue which adhered to the agarose on which the tissue was embedded. Dishes were filled with room temperature mammalian phosphate buffer and transferred to the inverted microscope to perform the measurements.

For nerve FBR capsules and epineurium measurements, implanted conduits were extracted from rats 3 months post-implantation with regenerated sciatic nerves still within them. Under a dissection microscope, fibrotic tissue covering the outside of conduits was removed. A cut was done along the length of the conduit, and the regenerated nerve fragment within was carefully removed. The nerve fragment was embedded on its side on a shallow bed of 4% w/w low melting point agarose (Sigma), and submerged in mammalian physiological saline. Finally, AFM measurements of the side of the nerve were carried out.

#### Material preparation

The stiffness of polyacrylamide hydrogel and silicone rubber implants and substrates was checked for every manufactured batch by AFM. Implants and substrates were all cleaned by immersion in PBS overnight. These were then transferred to 35 mm plastic dishes and fixed in place using a small amount of vaseline petroleum jelly (28908.290, VWR). Dishes were transferred to the inverted microscope to perform AFM measurements.

#### Indentation experiments

Petri dishes containing the samples to be analysed were placed on a motorised xy stage, which allowed movement of the sample relative to the AFM cantilever. A CCD camera (The Imaging Source GmbH) was used to image and track the position of the cantilever above the sample. This setup was used to locate and define an area of interest on the sample on which AFM measurements were taken. A custom python script broke down this area into 20 x 20 μm squares, inside which a single measurement was taken. The motorised stage was moved as measurements were taken to perform a raster scan of the area of interest.

For each elasticity measurement, the cantilever probe was lowered onto the surface of the sample at a speed of 10 μm/s. Upon contact and indentation of the sample, the probe continued to be lowered until a force of 10 nN was reached (usually equivalent to an indentation depth δ of 1 to 5 μm). The probe was then retracted, the sample moved, and a measurement repeated at a different location.

The force-distance measurements taken by the AFM were translated into elasticity values using the Hertz model^61^ using a previously described custom^62^ MATLAB (Mathworks, R2008a) script for every indentation of the sample. The cantilever and polystyrene bead probe on the sample was modelled as a sphere and a half space, and used to calculate the apparent reduced elastic modulus *K*.

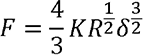

With *F* being the force applied and R the radius of the polystyrene bead. Elasticity was calculated at an indentation depth δ of 2 μm. The reduced elastic modulus may be further transformed into other elastic moduli, including Young’s modulus (*E*)^62^ and Shear modulus (*G*). A Poisson ratio ν of polyacrylamide was set to 0.48^63^, while for PDMS a value of 0.499 was used^64^.

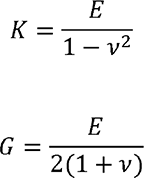

## Acknowledgements

The authors would like to thank Jessica Kwok for technical support and advice. Part of the RNA-Seq work was performed with the Genomics and Transcriptomics Core, which is funded by the UK Medical Research Council (MRC) Metabolic Disease Unit (MRC_MC_UU_12012/5) and a Wellcome Trust Major Award (208363/Z/17/Z), and guidance from Marcella Ma, whom the authors wish to thank. This work was furthermore supported by the UK Medical Research Council (MRC) and the Sackler Foundation (doctoral training grant RG70550 to ACL), the UK Wellcome Trust (Translational Medicine and Therapeutics PhD Programme Fellowship 109511/Z/15/Z to DGB), the Centre for Trophoblast Research (MP and RSH), the Whitaker International Scholars Program (ALR), the European Commission’s Horizon 2020 (Marie Sklodowska-Curie Fellowship 797506 to ALR), the Bertarelli Foundation (SPL), the European Research Council (Consolidator Award 772426 to KF), the UK Biotechnology and Biological Sciences Research Council (Research Grant BB/N006402/1 to KF), the UK Medical Research Council (Career Development Award G1100312/1 to KF), and the Alexander von Humboldt Foundation (Alexander von Humboldt Professorship to KF).

## Supplementary Material

**Supplementary Figure 1.**
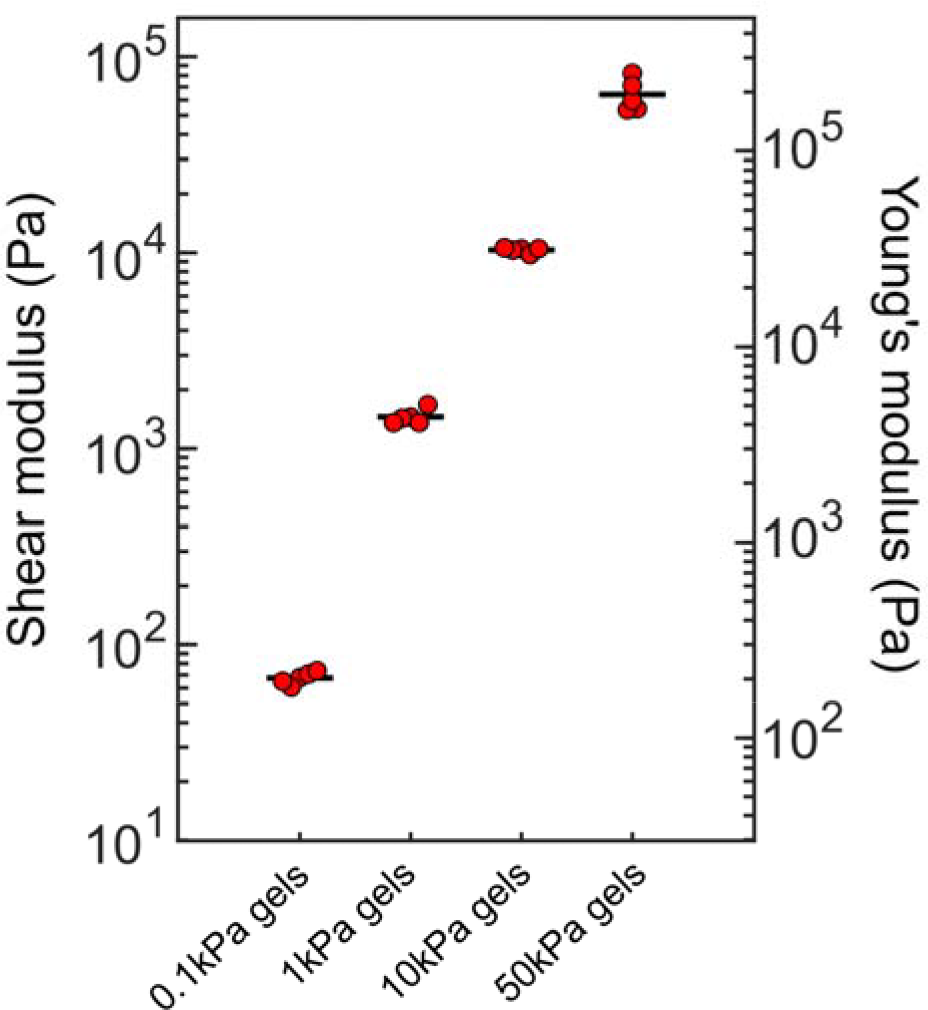
Plot of AFM-based stiffness measurements of polyacrylamide substrates used in *in vitro* experiments (Fig. 1-2). N=5 gels per condition. Values per gel represented by red circles; bars represent mean value. Stiffness values given in shear modulus (G) and Young’s modulus (E), based on a Poisson ratio of polyacrylamide gels of 0.48 ^63^.

**Supplementary Figure 2.**
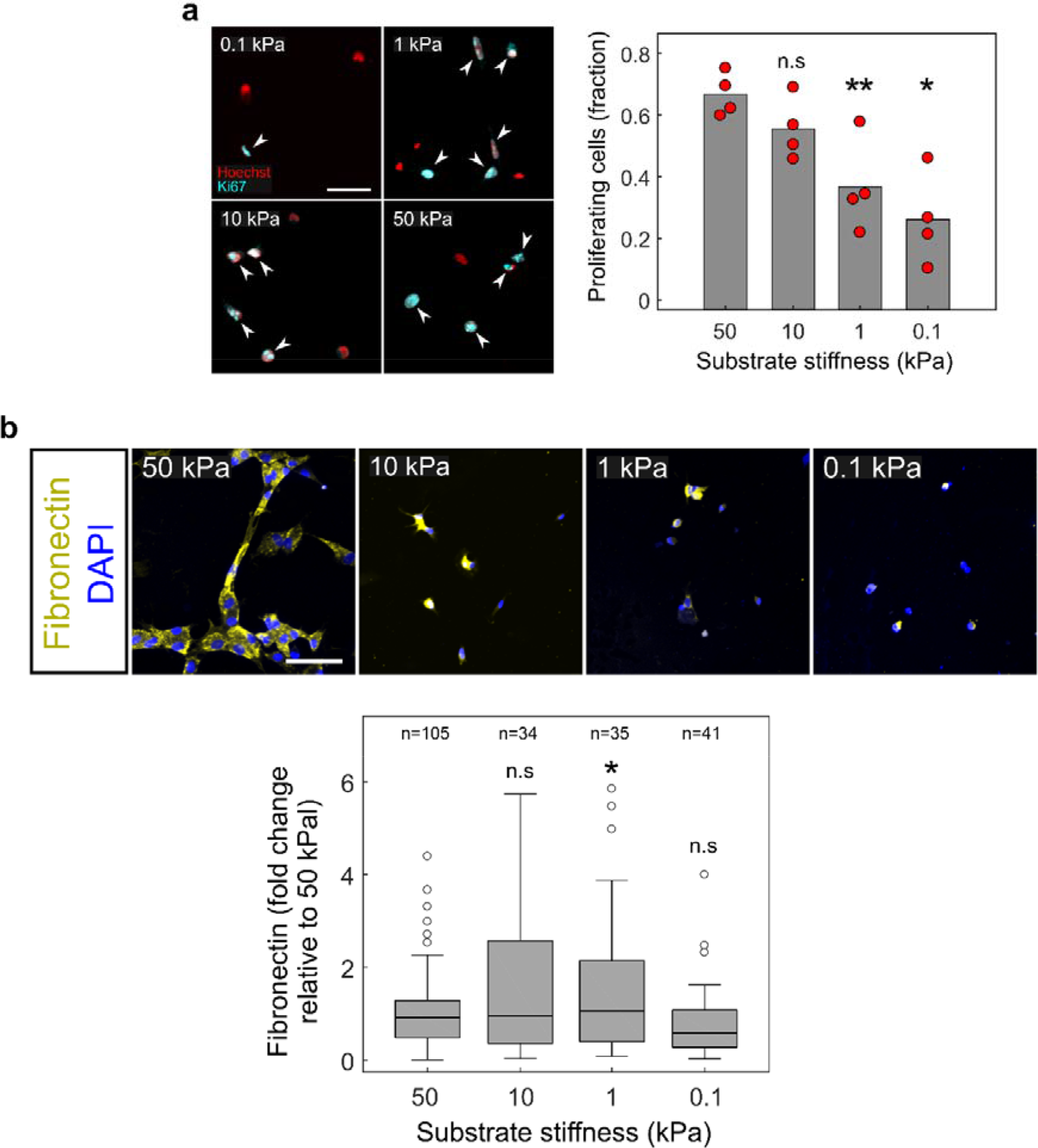
Fibroblast proliferation and fibronectin production as a function of substrate stiffness. **a,** Z-stack confocal images and quantification of nerve fibroblasts at 6 DIV cultured on polyacrylamide substrates of various stiffness (50, 10, 1 and 0.1 kPa shear modulus), stained for the proliferation marker Ki67, and the nuclear marker DAPI. Cells with overlapping DAPI and Ki67 stain (white arrowheads) are considered to be proliferating. Scale bar: 50 μm. N=4 cultures. Plot represents values for individual cultures (red circles) and mean of all cultures (bar). **b,** Z-stack confocal images and quantification (box plots) of nerve macrophages stained for the extracellular protein fibronectin, as well as nuclei (DAPI). Scale bar: 60 μm. n=number of cells. Box plot characteristics described in Methods. All statistical comparisons done via one-way ANOVA followed by Dunnett’s multiple comparisons test comparing to the 50 kPa condition. *, p<0.05; **, p<0.01, n.s, not significant (p>0.05).

**Supplementary Figure 3.**
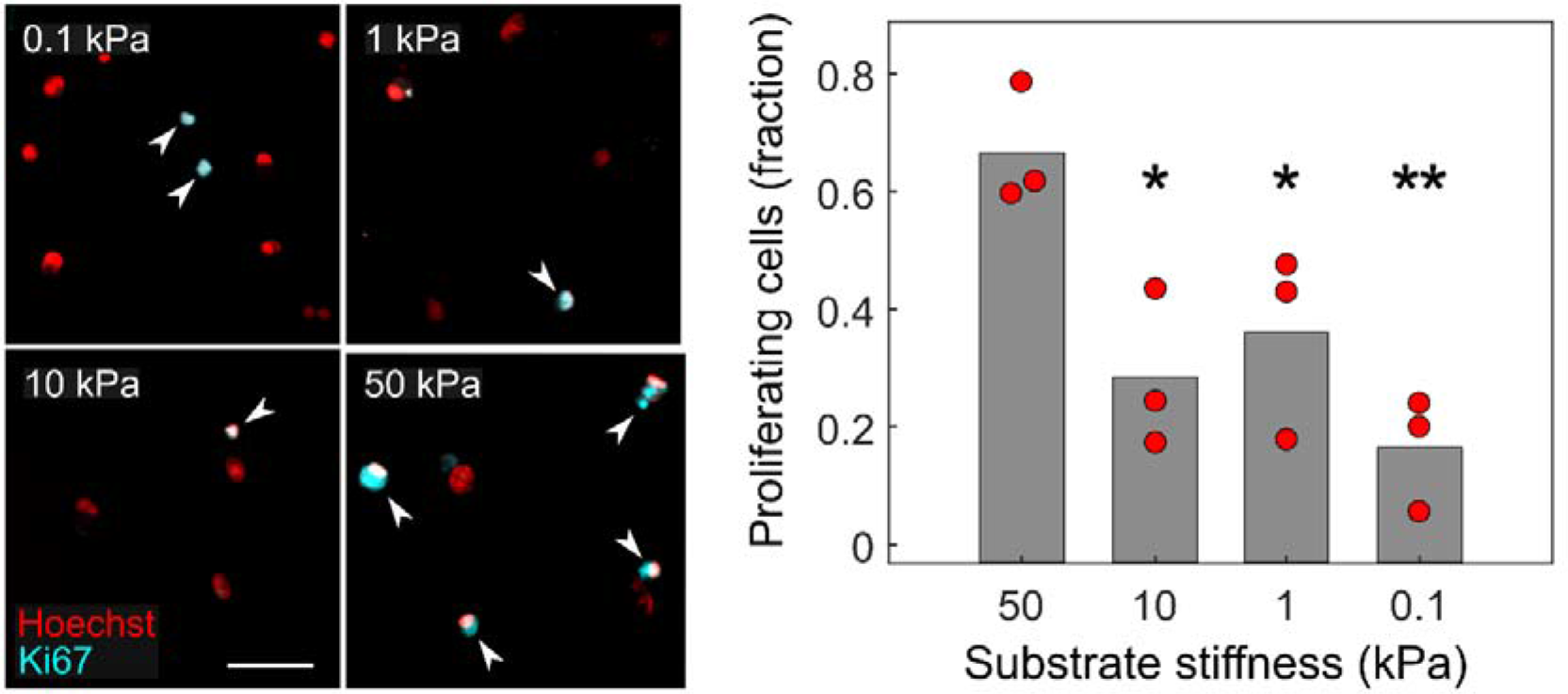
Macrophage proliferation as a function of substrate stiffness. Immunostaining and quantification of proliferating macrophages, stained for the marker Ki67, as well as the nuclear marker DAPI. Cells with overlapping DAPI and Ki67 stain (white arrowheads) are considered proliferating. Scale bar: 50 μm. N=3 cultures. Plot represents values for individual cultures (red circles) and mean of all cultures (bar). All statistical comparisons done via one-way ANOVA followed by Dunnett’s multiple comparisons test comparing to the 50 kPa condition. **, p<0.01; ***, p<0.001; n.s, not significant (p>0.05).

**Supplementary Figure 4.**
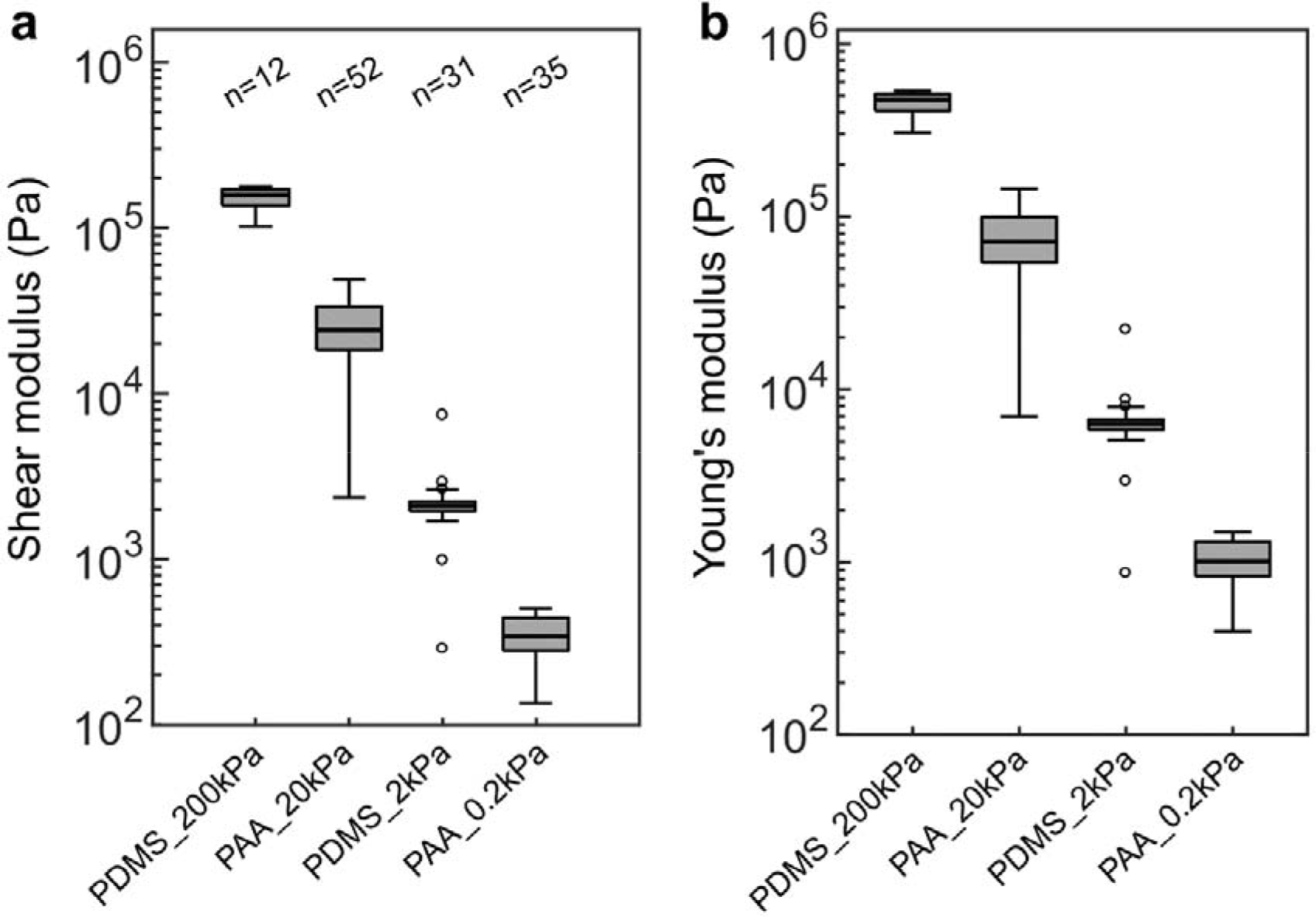
Stiffness measurements of materials used in *in vivo* experiments. Stiffness values given in terms of (**a**) shear modulus (*G*) and (**b**) Young’s modulus (*E*), based on a Poisson ratio of 0.48 for polyacrylamide gels^63^ and of 0.499 for silicone^64^. . n=number of measurements. Box plot characteristics described in Methods.

**Supplementary Figure 5.**
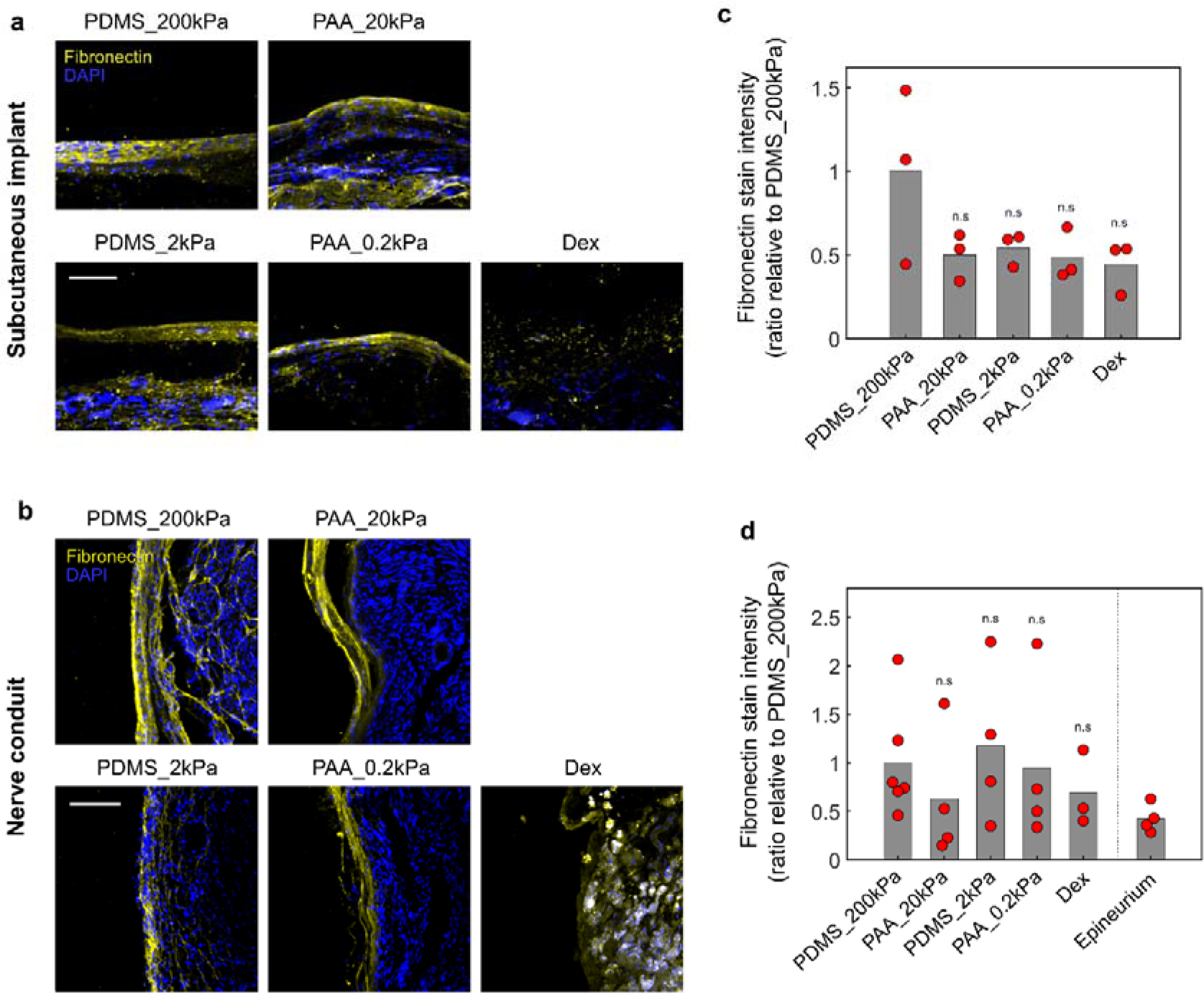
a,c, Fibronectin production *in vivo* is independent of implant stiffness. Z-stack confocal images of the FBR capsule developed over 3 months in response to coated implants in (**a**) subcutaneous tissue and (**b**) sciatic nerve, stained for fibronectin (yellow) and nuclei (DAPI, blue). Scale bars: 100 μm. **c**,**d**, Quantification of fibronectin stain intensity (values represented by red circles; mean represented by bar). All statistical comparisons carried out via one-way ANOVA followed by Dunnett’s multiple comparisons test comparing groups to the PDMS_200kPa control condition. Epineurium condition not included in statistical analysis. n.s, not significant (p>0.05).

**Supplementary Figure 6.**
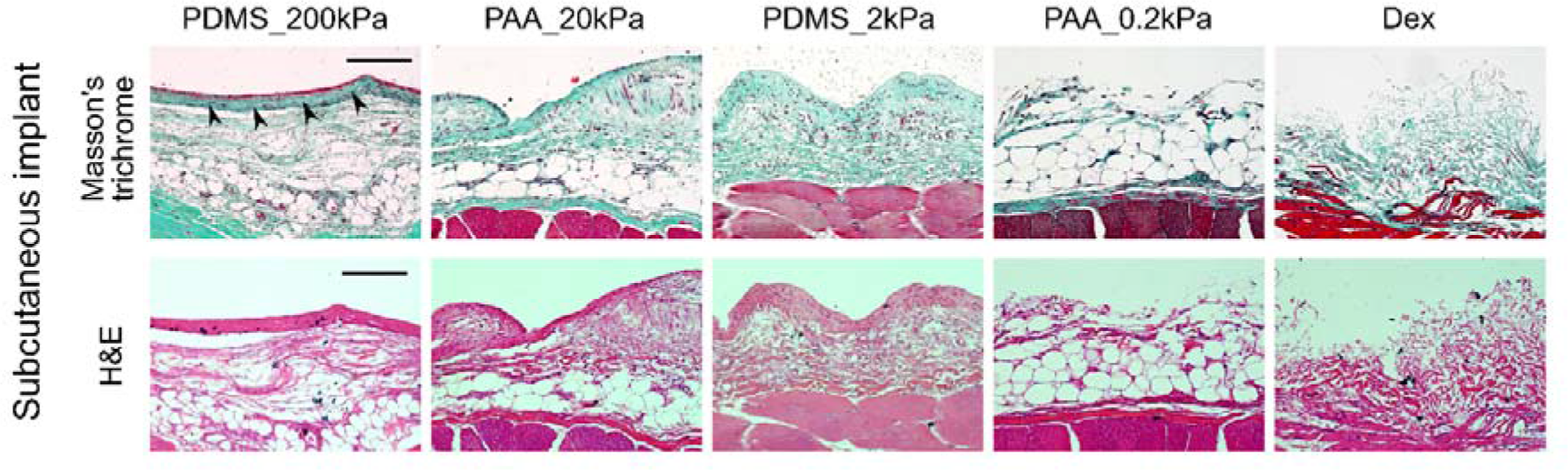
Masson’s trichrome (MT) and H&E stained sections of FBR tissue developed around subcutaneous implants after 3 months of implantation. Muscle and fascia visible at the bottom of images (red mass and blue collagen layer directly above it in MT stain). Stains reveal differences in tissue structure and composition of FBR capsules. Capsules around PDMS_200kPa are formed by a collagen-dense layer (black arrowheads), which is absent in all other conditions. PDMS_200kPa also presents a thicker layer of connective tissue separating the implant from muscle and fascia compared to all other conditions, indicative of a more intense FBR. Scale bar: 150 μm.

**Supplementary Figure 7.**
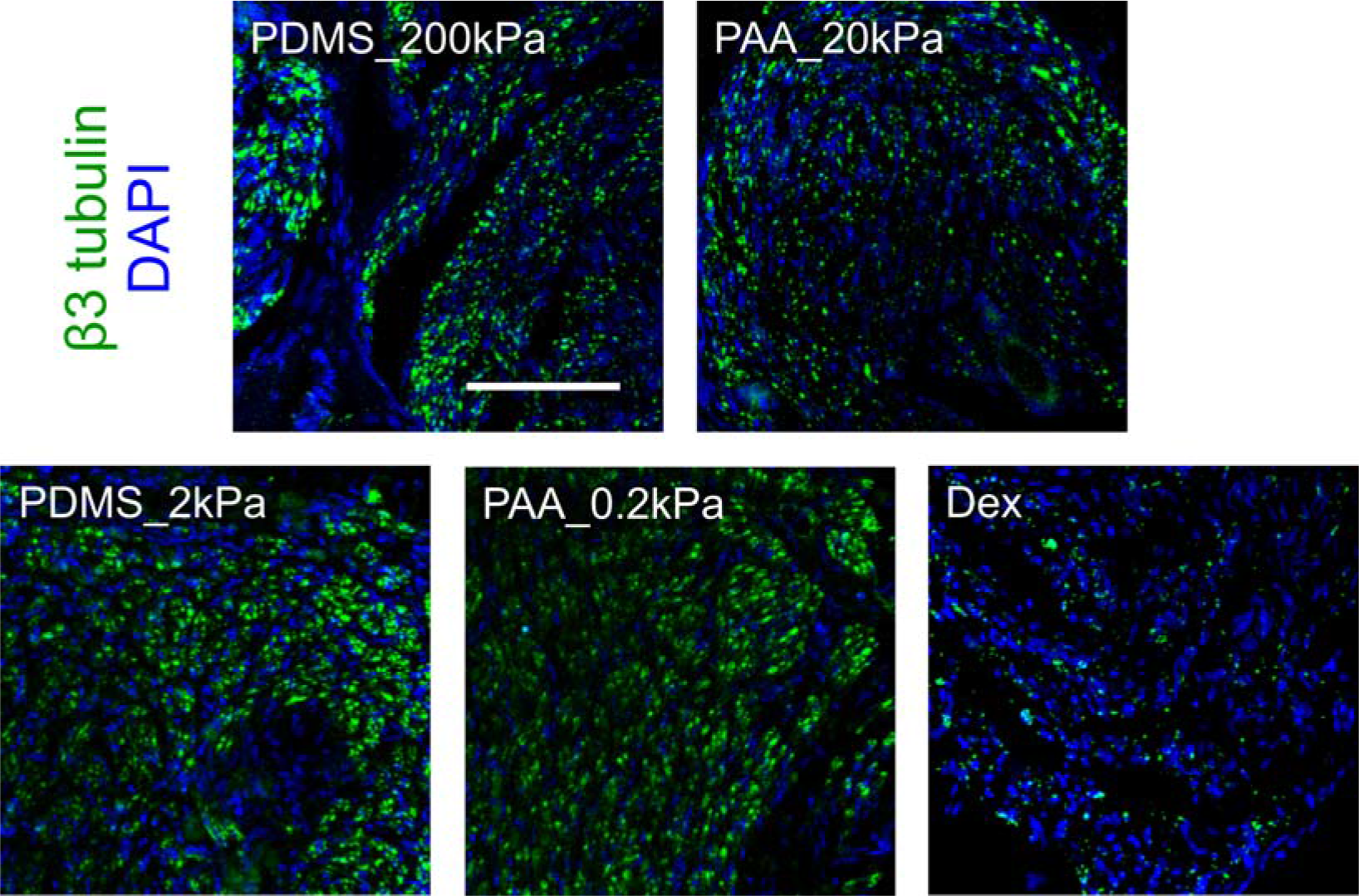
Dexamethasone impedes nerve regeneration. Z-stack confocal images of sciatic nerves exposed to implants of controlled stiffness (cross- sections), 3 months post-implantation. Cross-sections were taken 5 mm downstream of the point where the nerve was transected and the conduit was implanted. Axons stained in green (β3 tubulin), nuclei in blue (DAPI). Axon stains show a large decrease in axon density in nerves exposed to dexamethasone (Dex), while axon density is similar in all conditions with materials of controlled stiffness. Quantification of axon density in implanted nerves was carried out at this point of the nerve, downstream of the implantation site, and its quantification is shown in Figure 4e. Scale bar: 100 μm.

**Supplementary Figure 8.**
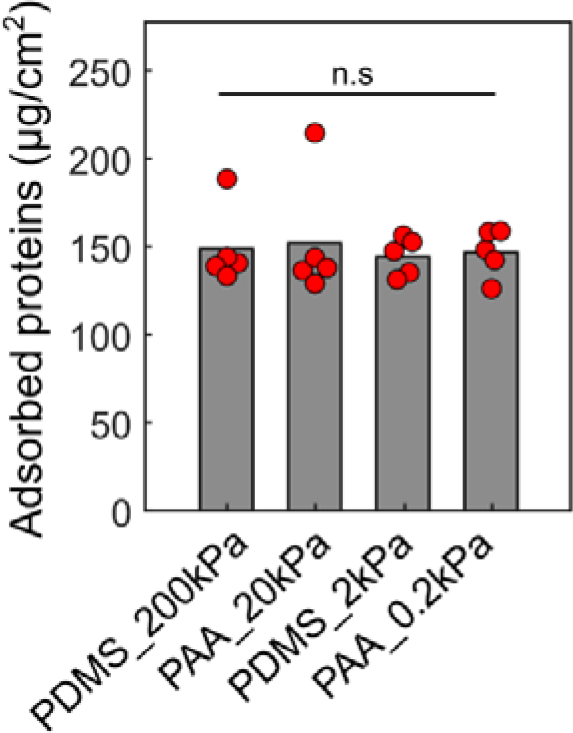
Protein adsorption assay. *In vitro* protein adsorption to materials of controlled stiffness used in *in vivo* implantation, following two-hour incubation in serum. N=5 samples of material per condition. Values per gel represented by red circles; bars represent mean value. No difference was observed in protein adsorption between all four materials. Statistical comparison done using one-way ANOVA (*p* = 0.954). n.s, not significant.

**Supplementary Figure 9.**
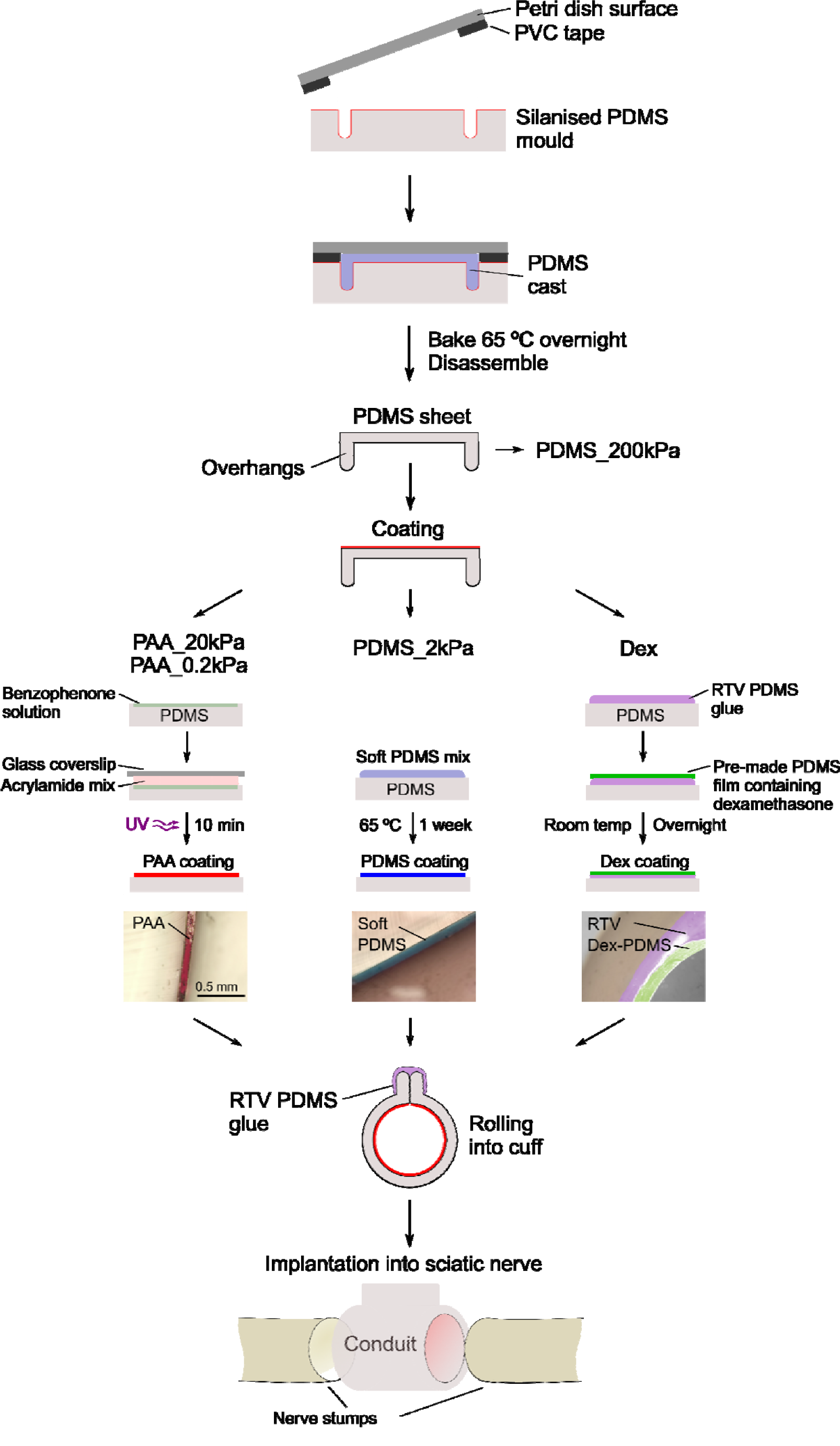
Nerve conduit fabrication protocol. Sheets of PDMS with overhangs are produced on a chamber formed by a silanized PDMS mould, petri dish and PVC tape. The resulting PDMS sheets were then coated on one of their surfaces with a material of specific stiffness, or kept uncoated for the stiff PDMS condition. **a.** For PAA hydrogel coatings, the PDMS surface was functionalised with benzophenone, and hydrogels were crosslinked under UV light. Red colouring has been added to PAA hydrogels in the image for easier visualisation. **b.** For soft silicone coatings, a drop of soft silicone mix was spread over the surface of the PDMS sheet, and baked at 65 °C for one week. Blue colouring has been added to the soft silicone in the image for easier visualisation **c.** For coatings containing dexamethasone, a small drop of RTV PDMS glue was spread over the surface of the PDMS sheet, and a thin sheet of PDMS containing dexamethasone laid down on top. The glue was allowed to set overnight. RTV glue and dex-containing PDMS are false coloured in image for easier visualisation. PDMS sheets were then rolled into conduits, and sealed to keep their shape using RTV PDMS glue applied to the overhangs. After allowing the glue to set overnight, conduits were UV sterilised and implanted into rat sciatic nerves.

**Supplementary Table 1.**
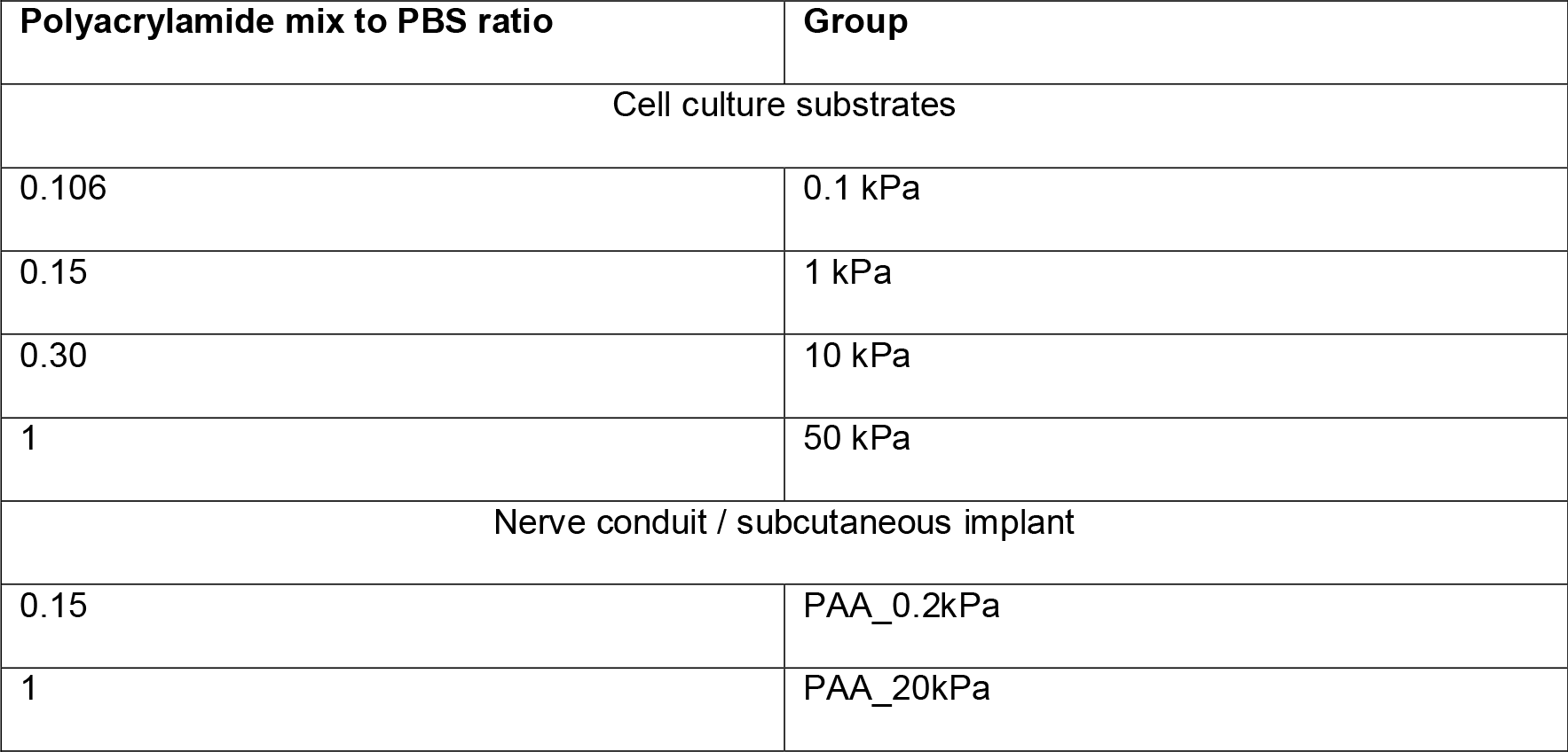
PAA protocols. Polyacrylamide hydrogel mixes used to produce the substrates and coatings of various stiffnesses. Note the polyacrylamide mixes used for cell culture experiments and for implants are slightly different (see Methods section).

| Polyacrylamide mix to PBS ratio | Group |
| --- | --- |
| Cell culture substrates |  |
| 0.106 | 0.1 kPa |
| 0.15 | 1 kPa |
| 0.30 | 10 kPa |
| 1 | 50 kPa |
| Nerve conduit / subcutaneous implant |  |
| 0.15 | PAA_0.2kPa |
| 1 | PAA_20kPa |

**Supplementary Table 2.** Antibodies used in immunocytochemistry and immunohistochemistry experiments.

| Antibody | Dilution | Source | Code |
| --- | --- | --- | --- |
| **Primary antibodies** |  |  |  |
| Rabbit anti-beta Tubulin | 1:200 | Abcam | ab6046 |
| Rabbit anti-Collagen I | 1:500 | Abcam | ab34710 |
| Rabbit anti-Fibronectin | 1:100 | Abcam | ab2413 |
| Rabbit anti-Ki67 | 1:100 | Abcam | ab15580 |
| Mouse anti-Vinculin | 1:100 | Abcam | ab18058 |
| Mouse anti-CD68 | 1:100 | Serotec | MCA341R |
| Mouse anti-αSMA | 1:100 | Abcam | ab7817 |
| Guinea pig anti-YAP1 | 1:500 | Abcam | ab195987 |
| Phalloidin-Alexa fluor 488 | 1:40 | ThermoFisher | a12379 |
| **Secondary antibodies** |  |  |  |
| Alexa Fluor donkey anti-mouse 555 | 1:1,000 | Invitrogen | ab150106 |
| Alexa Fluor donkey anti-rabbit 488 | 1:1,000 | Invitrogen | ab150073 |
| Alexa Fluor donkey anti-mouse 488 | 1:400 | Abcam | ab150185 |
| Hoechst-33342 nuclear stain | 1:10,000 | Sigma | 14533 |

## Notes

### Competing Interest Statement

The authors have declared no competing interest.

### Summary of Updates

We added several new exerimental data and data analyses.

